# Characterizing the portability of RecT-mediated oligonucleotide recombination

**DOI:** 10.1101/2020.04.14.041095

**Authors:** Gabriel T. Filsinger, Timothy M. Wannier, Felix B. Pedersen, Isaac D. Lutz, Julie Zhang, Devon A. Stork, Anik Debnath, Kevin Gozzi, Helene Kuchwara, Verena Volf, Stan Wang, Xavier Rios, Christopher J. Gregg, Marc J. Lajoie, Seth L. Shipman, John Aach, Michael T. Laub, George M. Church

## Abstract

Bacterial genome editing methods are used to engineer strains for biotechnology and fundamental research. Homologous recombination (HR) is the most versatile method of genome editing, but traditional techniques using endogenous RecA-mediated pathways are inefficient and laborious. Phage encoded RecT proteins can improve HR over 1000-fold, but these proteins have limited portability between species. Using *Escherichia coli*, *Lactococcus lactis, Mycobacterium smegmatis, Lactobacillus rhamnosus*, and *Caulobacter crescentus* we investigated the host-limited functionality of RecTs. We find that these proteins specifically recognize the 7 C-terminal amino acids of the bacterial single-stranded DNA-binding protein (SSB), and are portable between species only if compatibility with this host domain is maintained. Furthermore, in some species, we find that co-expressing otherwise incompatible RecTs with a paired bacterial SSB is sufficient to establish functionality. Finally, we demonstrate that high-efficiency HR surpasses the mutational capacity of more widely used error-prone methods for genome diversification, and can be used to identify exceptional phenotypes inaccessible through sequential nucleotide conversions.

## Main Text

Genome modification is crucial to the study of microbes, and is used to investigate the effects of genetic variation in genes and regulatory sequences^1^, to explore the properties of human commensals and pathogens^2,3,4,5^, and to engineer strains for biotechnology^6,7^. Homologous recombination (HR) is among the most versatile methods of genome editing, and relies on flanking homology arms to incorporate DNA into a targeted locus. It has no target site restrictions and in principle any type of genetic modification can be introduced. However, the endogenous pathway of RecA-mediated HR is often inefficient and requires long homology arms, making the introduction of genetic changes unreliable and limiting the generation of genomic libraries. Cas9, in contrast to its effects in mammalian cells, does not greatly improve the efficiency of HR in most prokaryotes^8^. Instead, since dsDNA breaks are often lethal, Cas9 is used as a tool to cut and select against non-edited cells^8^. High-throughput genome engineering in prokaryotes currently relies on overexpression of phage RecT proteins that integrate singlestranded oligonucleotide donors without nicking or breaking the genome (referred to as recombineering)^9,10,11,12^. Unfortunately, these proteins do not function broadly, and have only been established in a small number of species.

The Red operon from Enterobacteria phage λ is the source of one of the most widely used RecT proteins, λ-Red β (Redβ). Redβ, along with the rest of the λ-Red operon, enables a suite of technologies including gene knockouts using PCR products^10^, multiplex genome editing^13,6^, and state-of-the-art methods for synthetic genome assembly^14,15^. It also provides one of the most rigorous and comprehensive methods for generating libraries of genetic varients^1,3^. Rather than cloning and screening mutant genes on plasmids^16^, or generating nucleotide variants through random mutagenesis^17^, recombineering can be used to precisely mutate genes in their endogenous context. Redβ, however, only operates in a small set of phylogenetically related species (*E. coli, Salmonella enterica* and *Citrobacter freundii*, among others)^18^. The key features influencing the host-tropism exhibited by Redβ and other RecT proteins are not well understood^19^. In this work, we aimed to characterize the key features limiting the portability of Redβ and other RecT protein homologs and extend the host-range of recombineering.

The most reliable strategy for establishing recombineering in species distantly related to *E. coli* has been to screen RecT proteins from phages or prophages that infect the host of interest^20,21^. We therefore hypothesized that phage RecT proteins may require a specific interaction with one or more bacterial proteins, limiting their functionality to species where this host-protein interaction is maintained. A key mechanistic step during recombineering is annealing of ssDNA to the genome at the lagging strand of the replication fork^19^. We thus focused our efforts on host proteins which reside or consistently interact with DNA on the lagging strand. Recently, we identified a binding interaction between Redβ and a C-terminal peptide of *E. coli* SSB (singlestranded DNA-binding protein)^22^. Phage SSBs have been shown to play important roles in phage replication and recombination pathways^23^, and SSBs have been used to increase recombineering efficiency in *E. coli*^24^. In this work, we explored whether a specific interaction between phage RecTs and bacterial SSBs might generally influence the portability of recombineering methods.

Here, we provide evidence that a specific interaction between phage RecT proteins and the host SSB limits their portability. We find the majority of this interaction relies on recognition of SSB’s 7 C-terminal amino acids. We also find that, in some species, supplying an exogenous bacterial SSB can make up for a lack of host-compatibility, allowing RecT proteins to function in previously recalcitrant species. We then demonstrate the potential for high-efficiency recombineering in new species by optimizing recombineering in *L. lactis*. Specifically, we find that oligonucleotide recombination can be used to navigate genotypic landscapes that contain extensive epistatic effects, and are inaccessible through error-prone methods which primarily generate single-nucleotide conversions.

## Results

### SSB is a key mediator of RecT host specificity

To understand the host-tropism displayed by RecT’s, we started by developing a simplified *in-vitro* model of oligonucleotide annealing that includes bacterial SSBs, a key host protein that coats ssDNA at the replication fork^25^. We first tested whether two 90bp oligos could anneal if they were pre-coated with SSB. We purified SSBs from *E. coli* (gram-negative), where most recombineering work has been performed, and *L. lactis* (gram-positive), a lactic acid bacterium phylogenetically distantly related to *E. coli*. Using fluorescence quenching to measure annealing, we found that while the free oligos annealed together slowly (Fig 1a,b), both EcSSB and LlSSB completely inhibited oligonucleotide annealing. We next tested capacity of a phage RecT protein to overcome this SSB-mediated inhibition of annealing. We thus purified Redβ, which is not broadly portable, but mediates efficient oligonucleotide annealing in *E. coli*. We found that adding Redβ overcame the inhibitory effect of EcSSB but not LlSSB, rapidly annealing the EcSSB-coated two oligos together (Fig 1a,c). These preliminary results gave us an indication that while bacterial SSBs inhibit oligonucleotide annealing *in vitro*, RecTs overcome the inhibitory effect in an SSB-specific manner.

**Fig. 1.**
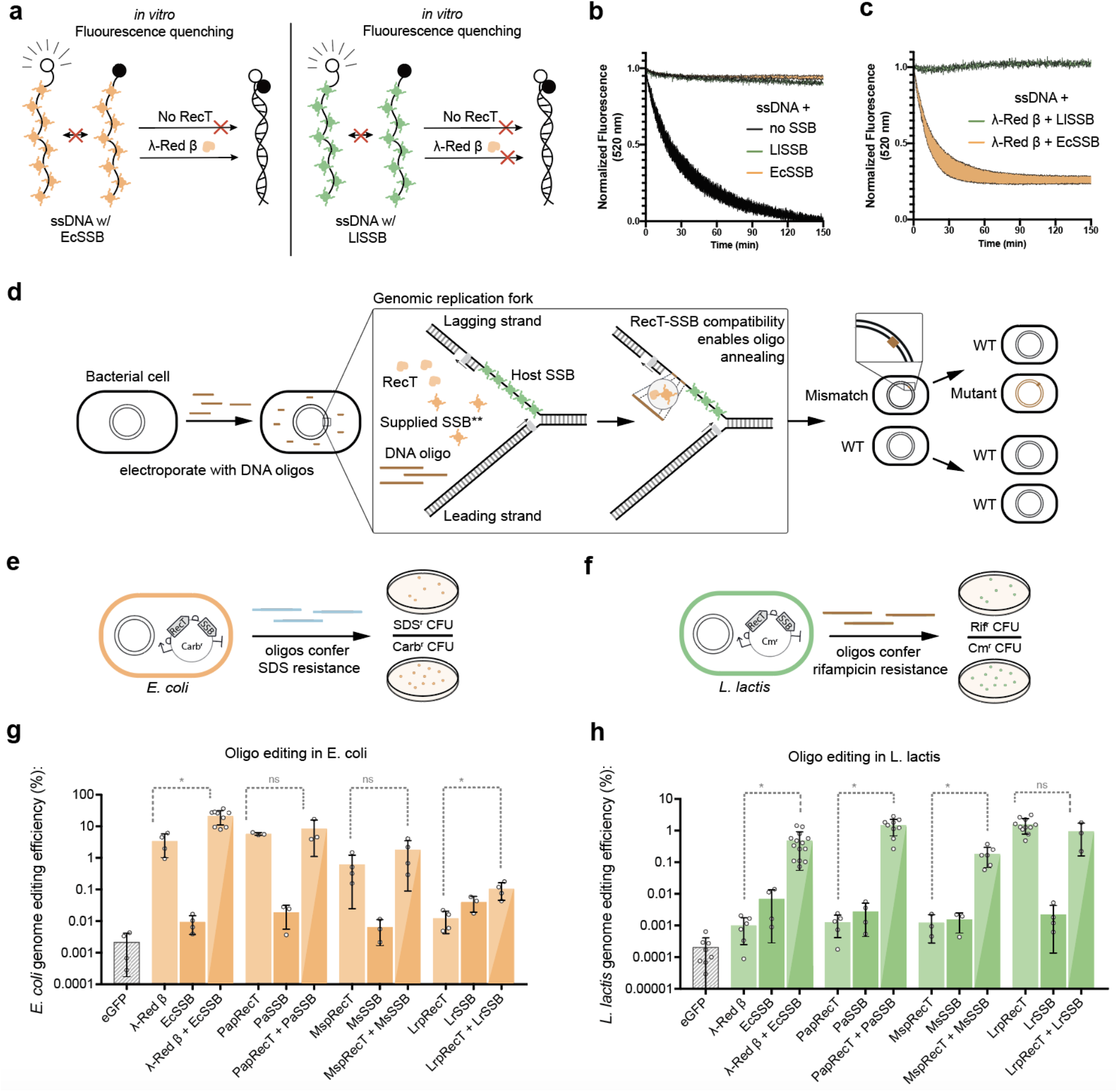
(a) In-vitro model of ssDNA annealing inhibition by EcSSB or LlSSB, and ability of λ-Red β to overcome annealing inhibition by EcSSB. (b) ssDNA annealing without SSB, precoated with EcSSB, or pre-coated with LlSSB. Shaded area represents the SEM of at least 2 replicates (c) ssDNA annealing in the presence of λ-Red β when pre-coated with EcSSB or LlSSB. Shaded area represents the SEM of at least 2 replicates (d) Model for RecT-mediated editing in the presence of SSB. An interaction between RecT and the host SSB enables oligo annealing to the lagging strand of the replication fork. **Co-expressing an exogenous SSB that is compatible with a particular RecT variant can in some species enable efficient homologous genome editing even if host compatibility does not exist. (e),(f), Calculation of editing efficiency in *L. lactis* and *E. coli* is performed by introducing antibiotic resistance mutations into the genome using synthetic oligos, and then measuring the ratio of resistant cells to total cells (g),(h), A comparison of the efficiency of editing in *L. lactis* and *E. coli* after the expression of either RecTs, SSBs, or “cognate pairs” (as described in the text). Lighter colors represent RecT expression, while darker colors represent SSB expression. Mixed light/dark bars represent coexpressed RecTs-SSB pairs.

To validate this result *in vivo*, we developed an assay to measure the portability of RecT proteins. We selected four variants known to enable high genome editing efficiency (Redβ and PapRecT in *E. coli*, LrpRecT in *L. lactis*, and MspRecT in *M. smegmatis*), and tested codon optimized versions of all four in both *E. coli* (gram-negative), and *L. lactis* (gram-positive). We measured genome editing efficiency by introducing oligos encoding known antibiotic resistance mutations, and compared the antibiotic resistant cell counts to the total number of viable cells in the population (Fig 1e, f). In *E. coli*, Redβ and PapRecT functioned well, and improved oligo incorporation 1600-fold and 2700-fold respectively, while MspRecT (290-fold improvement) and LrpRecT (5.6-fold improvement) were less effective (Fig 1g). In *L. lactis*, LrpRecT was the only functional homolog, and improved oligo incorporation 7,700-fold, while the three other RecT variants were nearly non-functional, improving oligo incorporation less than 7-fold (Fig 1h). No RecT protein functioned well both in *E. coli* and *L. lactis*. This agrees with previous studies, which have found that RecT proteins are usually not portable between distantly related bacterial species^26^.

If interaction with the bacterial SSB is required for phage RecT functionality, one solution to establishing recombineering in a new species would be to replace the host SSB with one compatible with the chosen RecT. However, SSB proteins are essential, and mutations to SSB can result in severe growth defects^25^. We therefore evaluated if temporary overexpression of an exogenous SSB could supply the necessary requirements for recombineering and improve the activity of non-host compatible RecTs. We synthesized SSBs corresponding to each RecT protein (Note S1) and tested the activity of all four cognate RecT-SSB pairs in *L. lactis* and *E. coli*. Co-expression of a cognate bacterial SSB improved the genome editing efficiencies of all RecTs with low host-compatibility (Fig 1g, h). The best performing pairs, Redβ + EcSSB and PapRecT + PaSSB demonstrated 483-fold and 1,168-fold improved editing efficiencies over the RecT proteins alone in *L. lactis*, and still maintained high activity in *E. coli* (Fig 1g, h). In *E. coli*, co-expression with cognate SSBs also significantly reduced the toxicity of these two pairs (Fig S4). These results, especially in *L. lactis*, indicate that the presence of a cognate bacterial SSB can overcome the host incompatibility of RecT proteins if moved to new species.

### Compatibility between RecTs and SSBs is mediated by the 7 C-terminal amino acids of SSB

We next investigated which domains on SSB were involved in mediating the RecT protein interaction. A SSB domain-specific model for understanding RecT protein portability would be far more informative than previous models, which relied on phylogenetic relationships between the host organisms^26^. RecT proteins have been shown to function in species with SSBs with relatively divergent sequences. We were therefore interested in identifying conserved domains responsible for maintaining the RecT protein interaction. For example, while Redβ works well in *E. coli, Salmonella enterica*, and *Citrobacter freundii* which have SSBs with 88% identity, PapRecT works in *E. coli* and *Pseudomonas aeruginosa*, which have SSBs of only 59% identity. To investigate the specific residues involved, we used our genome editing assay in *L. lactis* and evaluated the effect of co-expressing RecT proteins with non-cognate or mutated SSBs.

The C-terminal tail of *E. coli* SSB is known to be the binding domain for host proteins involved in DNA replication and repair^25^, and a 9-amino-acid EcSSB C-terminal tail peptide was shown to bind to Redβ *in vitro*^22^. We therefore evaluated the importance of the SSB C-terminal tail by coexpressing Redβ in *L. lactis*, along with a version of EcSSB that had a 9-amino-acid C-terminal deletion (EcSSBΔ9) (Fig 2a). In *L. lactis*, the genome editing efficiency of Redβ with EcSSBΔ9 was 44-fold lower than Redβ with EcSSB, indicating a key role for the C-terminal domain in the SSB-mediated efficiency improvement (Fig 2c). Next, we co-expressed Redβ with the *L. lactis* SSB (LlSSB), which we expect to have little-to-no compatibility. Co-expression of Redβ with LlSSB performed similarly to Redβ with EcSSBΔ9, and improved genome editing efficiency 38.5-fold less than Redβ with EcSSB. We then co-expressed Redβ with chimeric versions of the LlSSB, where up to 9 amino acids of the LlSSB C-terminal tail were replaced with their corresponding residues from EcSSB. Swapping the last 7 C-terminal residues (LlSSB C7:EcSSB) improved editing rates to within 5.9-fold of Redβ with EcSSB, and swapping the last 8 C-terminal residues (LlSSB C8:EcSSB) improved editing rates within 2.6-fold of Redβ with EcSSB. These results support a model where Redβ specifically recognizes at minimum the 7 C-terminal acids of *E. coli* SSB, but not that of *L. lactis* SSB.

**Fig. 2.**
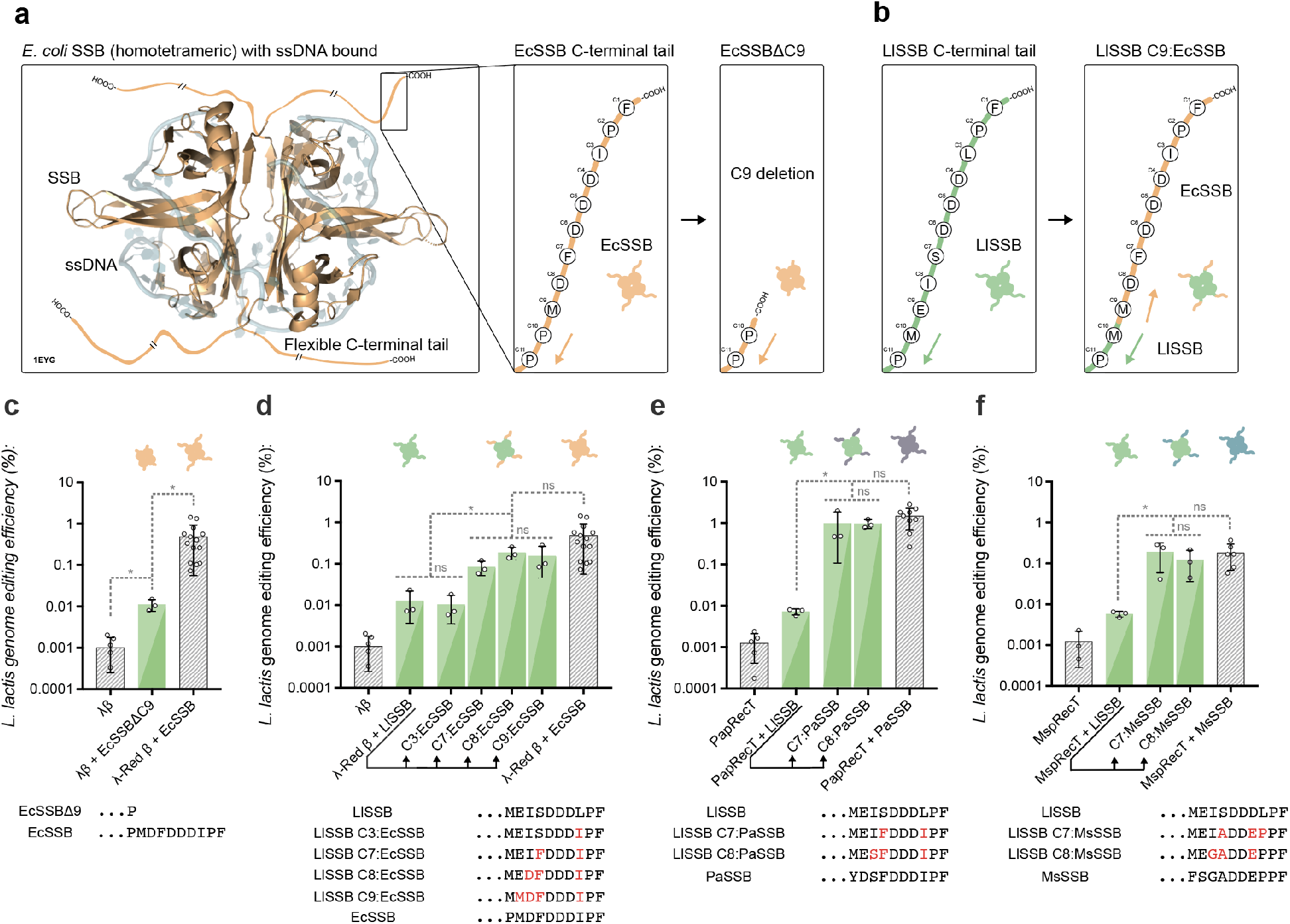
(a), A crystal structure of homotetrameric *E. coli* SSB bound to ssDNA (PDB-ID 1EYG)^37^. The amino acid sequence of the flexible C-terminal tail is diagramed in the right panel, along with the design of a 9AA C-terminal truncation to SSB. (b), The *L. lactis* SSB C-terminal tail is diagramed, along with an example of an SSB C-terminal tail replacement. In this case, the 9 C-terminal amino acids of the *L. lactis* SSB are replaced with the corresponding residues from *E. coli* SSB. The notation “LlSSB C9:EcSSB” is used as shorthand. (c), Editing efficiency in *L. lactis* of λ-Red β with a 9AA C-terminally truncated EcSSB mutant. (d), Editing efficiency in *L. lactis* of λ-Red β expressed with LlSSB, or mutants of LlSSB with C3, C7, C8, or C9 terminal residues replaced with the corresponding residues from EcSSB. (e, f) Editing efficiency in *L. lactis* of PapRecT (e) or MspRecT (f) expressed with LlSSB, or mutants of LlSSB with the C7 or C8 terminal residues replaced with the corresponding residues from the cognate SSB.

To evaluate if the SSB C-terminal 7 amino acids also affected the compatibility of the other two non-host compatible RecT proteins, we performed similar SSB-chimera experiments with PapRecT and MspRecT in *L. lactis*. The genome editing efficiency of PapRecT with the *L. lactis* SSB was 135-fold less than when using the cognate pair (Fig 2e). However, this defect was completely recovered when PapRecT was co-expressed with *L. lactis* SSB chimeras where either the last 7 or 8 C-terminal resides were replaced (LlSSB C7:PaSSB, LlSSB C8:PaSSB) (Fig 2e). For MspRecT, the genome editing efficiency with LlSSB was 33-fold lower than when using the cognate pair (Fig 2f). Again, the defect was completely recovered when MspRecT was co-expressed with *L. lactis* SSB chimeras where either the last 7 or 8 C-terminal resides were replaced (LlSSB C7:MsSSB, LlSSB C8:MsSSB) (Fig 2f). Since the chimeric LlSSBs greatly improved the functionality of non-host compatible RecT proteins, while the wild-type LlSSB did not, the RecT-SSB interaction seems to be both specific and relatively modular, with the 7 C-terminal amino acids acting as the critical interaction domain.

These results provide a molecular basis for the portability of RecT proteins between species which have host SSBs with a conserved C-terminal tail. Specifically, although the SSBs have only 59% identity, the *P. aeruginosa* and *E. Coli* SSBs have a perfectly conserved 7 amino acid C-terminal tail domain (Fig 3c), supporting the functionality of PapRecT in *E. coli*. Additionally, *E. coli, Salmonella enterica* and *Citrobacter freundii*^18^ SSBs all have a perfectly conserved 7 amino acid C-terminal tails, supporting the portability of Redβ between these species (Fig 3c).

**Fig. 3.**
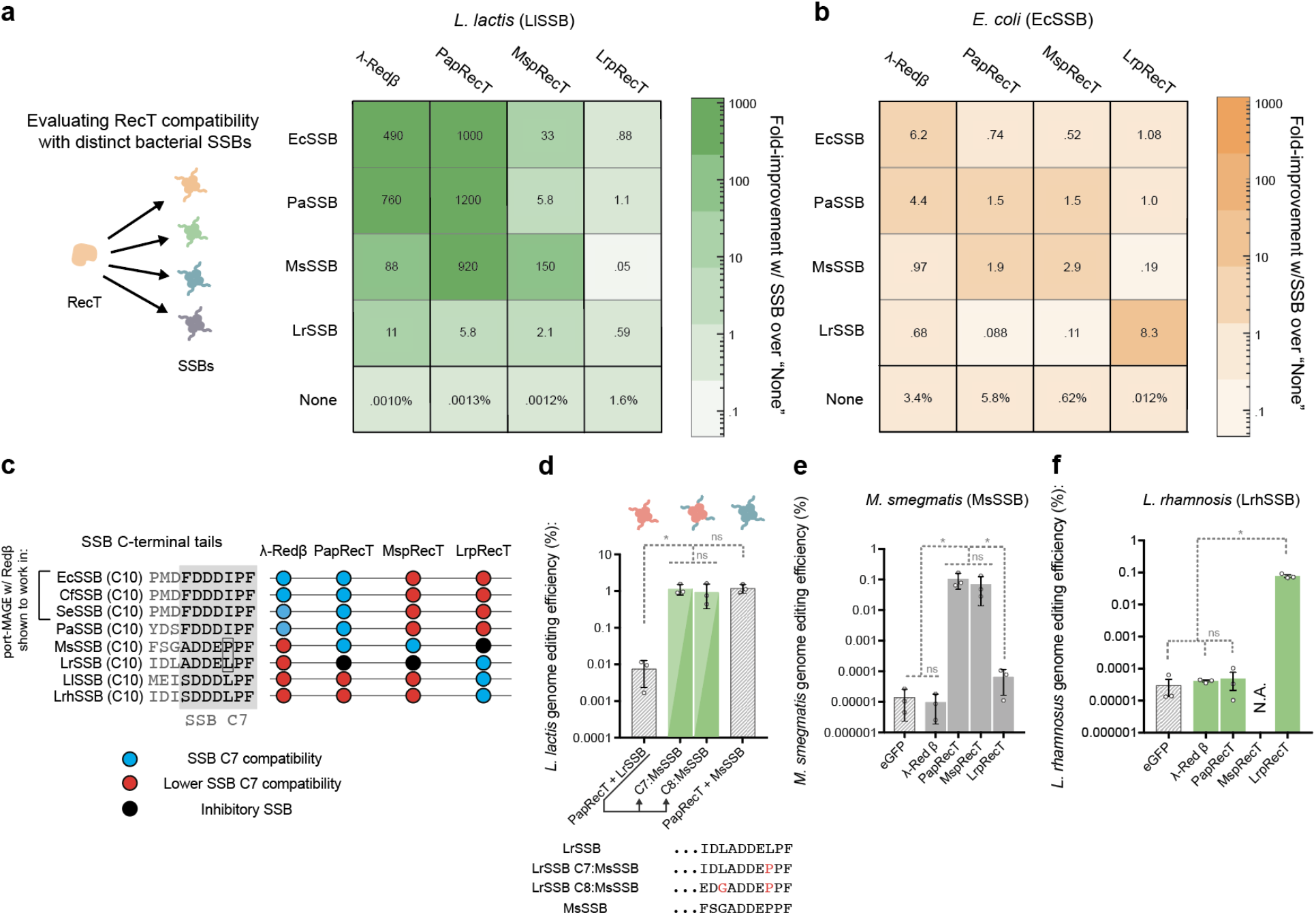
(a, b) Heat map showing the fold improvement in editing efficiency due to SSB coexpression in (a) *L. lactis* or (b) *E. coli* of RecT-SSB pairs as compared to the RecT alone. (c), C-terminal sequences of SSBs as well as RecT compatibility given “A” and “B”, for compatibility cutoffs given in Note S2. (d), Editing efficiency in *L. lactis* of PapRecT coexpressed with LrSSB, MsSSB, or mutants of LrSSB which had the C7 or C8 terminal residues replaced with the corresponding residues from the MsSSB. (e), Editing efficiency in *M. smegmatis* of λ-Red β, PapRecT, MspRecT, and LrpRecT. (f), Editing efficiency in *L. rhamnosus* of λ-Red β, PapRecT, MspRecT, and LrpRecT.

### Interactions between RecTs and SSBs are promiscuous and match basal RecT portability

Some RecT proteins are known to be portable between species which have distinct SSB C-terminal tails. To better characterize the network of RecT-SSB compatibility among the proteins analyzed here, we co-expressed all four RecTs with all four SSBs in both *E. coli* and *L. lactis* (Fig 3a, b, S3, S4). We found that the effects of PaSSB and EcSSB on RecT-mediated editing efficiency were relatively interchangeable, as might be expected since they share the same 7 amino acid C-terminal tail (Fig 3c). Interestingly, PapRecT displayed the characteristics of a more portable RecT protein, and showed compatibility with MsSSB and EcSSB/PaSSB, even though their 7AA C-terminal tail sequences are distinct (Fig 3a, c). Importantly, co-expressing PapRecT with LrSSB did not provide a substantial improvement in genome editing efficiency in *L. lactis*, even though the 7 C-terminal tail amino acids of LrSSB differ only by a single residue from MsSSB (Fig 3a, c).

To test if PapRecT specifically interacts with the C-terminal tail of MsSSB, we co-expressed PapRecT with a chimeric version of LrSSB, with either the C7 or C8 amino acids matching that of MsSSB (Fig 3d). The chimeric constructs demonstrated the same editing efficiency as PapRecT + MsSSB, showing that a single amino acid change was sufficient to enable compatibility between the proteins (Fig 3d). The compatibility of PapRecT with the distinct EcSSB/PaSSB and MsSSB tails but not the LrSSB tail affirms that while the SSB C-terminal tail has a critical role in the RecT-SSB interaction, there can be flexibility in the specific motif recognized.

We next evaluated if the interaction between PapRecT and MsSSB in *L. lactis* indicated that PapRecT would function in *M. smegmatis*, where MsSSB is natively expressed. We tested all four RecT’s in this species and indeed found that in *M. smegmatis* PapRecT enabled high efficiency editing, incorporating oligos at the same rate as MspRecT ^27^, while the other two RecT variants had much lower efficiency (Fig 3e).

Finally, we used our model for RecT to establish recombineering in *L. rhamnosus*, a well-studied probiotic used to treat a variety of illnesses including diarrhea and bacterial vaginosis. Although the *L. rhamnosus* SSB and *L. lactis* SSB only have 47% identity, they share identical SSB C-terminal tail 7 amino acids. Based on this we expected that LrpRecT (which functions in *L. lactis*) should be portable to *L. rhamnosus*, while the other RecT proteins would not be functional. We tested the 4 RecT proteins in this species and indeed found that LrpRecT incorporated oligonucleotides three orders of magnitude above the background level, while Redβ and PapRecT had negligible activity and MspRecT was toxic.

### RecT-SSB pairs can enable oligonucleotide recombination in organisms with no identified host-specific RecT

In *L. lactis*, the co-expression of PapRecT and Redβ with compatible SSB’s improved genome editing efficiency to a level comparable with the host-compatible LrpRecT. We hypothesized that for some species, RecT-SSB pairs could provide functional recombineering capacity even if no functional RecT protein had previously been identified. We therefore tested the two best-performing RecT-SSB pairs (Redβ + PaSSB, and PapRecT + PaSSB) for activity in *Caulobacter crescentus*, a model organism for studying cell cycle regulation, replication, and differentiation.

In *C. crescentus*, we did not detect any significant editing enhancement over the background with the RecT proteins alone, or PapRecT + PaSSB. As we previously observed compatibility between PapRecT and PaSSB, we believe additional factors must contribute to the incompatibility of this pair with *C. crescentus*. However, using Redβ + PaSSB, we observed a 15-fold improvement over Redβ alone (Fig 4a). After expression optimization (Fig S5) and evasion of mismatch repair, Redβ + PaSSB demonstrated 873-fold improved editing efficiency over the background level, which was 112-fold higher than Redβ alone (Fig 4b). These results indicate that while RecT-SSB pairs are not universally portable (Figure S6), the co-expression of a RecT protein with a compatible bacterial SSB will equal or surpass the editing efficiencies of RecT proteins alone.

**Fig. 4.**
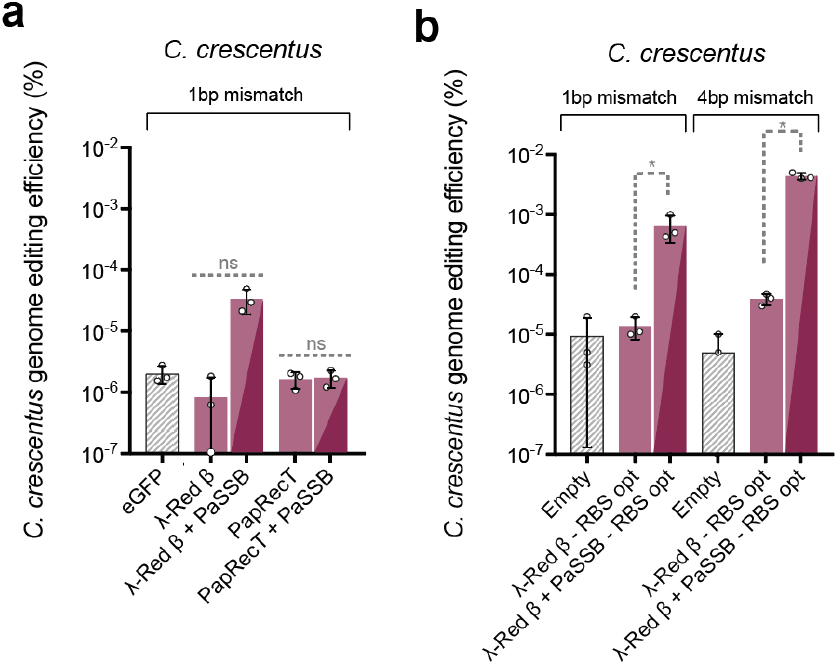
(a), Editing efficiency in *C. crescentus* of two RecT-SSB protein pairs, λ-Red β + PaSSB and PapRecT + PaSSB which had high genome editing efficiency in both *E. coli* and *L. lactis*. (b), Editing efficiency in *C. crescentus* of λ-Red β + PaSSB with ribosomal binding sites optimized for translation rate and using an oligo designed to evade mismatch repair.

### The utility of recombineering for generating targeted genomic libraries

In *E. coli*, one of the unique capabilities of recombineering is the ability to generate rationally designed or high-coverage genomic libraries. Although this technique (termed MAGE) has been used for a variety of applications including optimizing metabolic pathways^6^, protein evolution^28^, and saturation mutagenesis^1^, it has only been used in a limited capacity in other species. We used *L. lactis,* a microbe distantly related to *E. coli*, to demonstrate how mismatch repair evasion and oligonucleotide library design can be used to perform high-coverage genomic mutagenesis after a functional RecT protein has been identified.

To begin, we adapted our assay in *L. lactis* to allow the efficient incorporation of single, double, or triple nucleotide mutations, which are normally recognized and corrected by mismatch repair pathways. We used the cognate pair PapRecT and PaSSB, and co-expressed either the dominant negative mismatch repair protein MutL.E32K from *E. coli*^18^, or the host protein *L. lactis* MutL carrying the equivalent mutation (LlMutL.E33K, Fig S7). While MutL.E32K from *E. coli* was nonfunctional, co-expression of LlMutL.E33K enabled the efficient introduction of 1bp pair changes (Fig S8). Optimization of inducer and oligonucleotide concentrations further improved editing efficiency 26-fold (Fig S8).

Next, we used our optimized protocol to characterize the landscape of spectinomycin resistant variants at the ribosomal gene RpsE. This locus was previously studied in *E. coli* using an efficient error-prone mutagenesis method, and the genotypes sampled reflect the capabilities of other methods that generate nucleotide changes including hyper-mutator strains, and the amplification and cloning of genes using error-prone PCR. Since the crystal structure of RpsE bound to spectinomycin has long been known^29^, the main unknowns are the space of possible antibiotic resistance mutants, and the fitness of the different resistant mutants identified.

We started by targeting a 5-residue region near the spectinomycin binding pocket (*L. lactis* Lys 29), which is perfectly conserved between *L. lactis, E. coli* and *N. gonorrhea*, species for which spectinomycin has been used clinically^30^ (Fig 5a, Fig S15). We began by pooling 5 oligos, each targeted to single codon, to sample all possible single nucleotide conversions in the region (5×1NNK, 100 amino acids variants) (Fig 5a). We also synthesized an oligo pool targeting all 5 codons, and generated an estimated 96.7% of the possible combination mutants between the positions (5NNK, 3.2 million amino acid variants, Note S3) (Fig 5a).

**Fig. 5.**
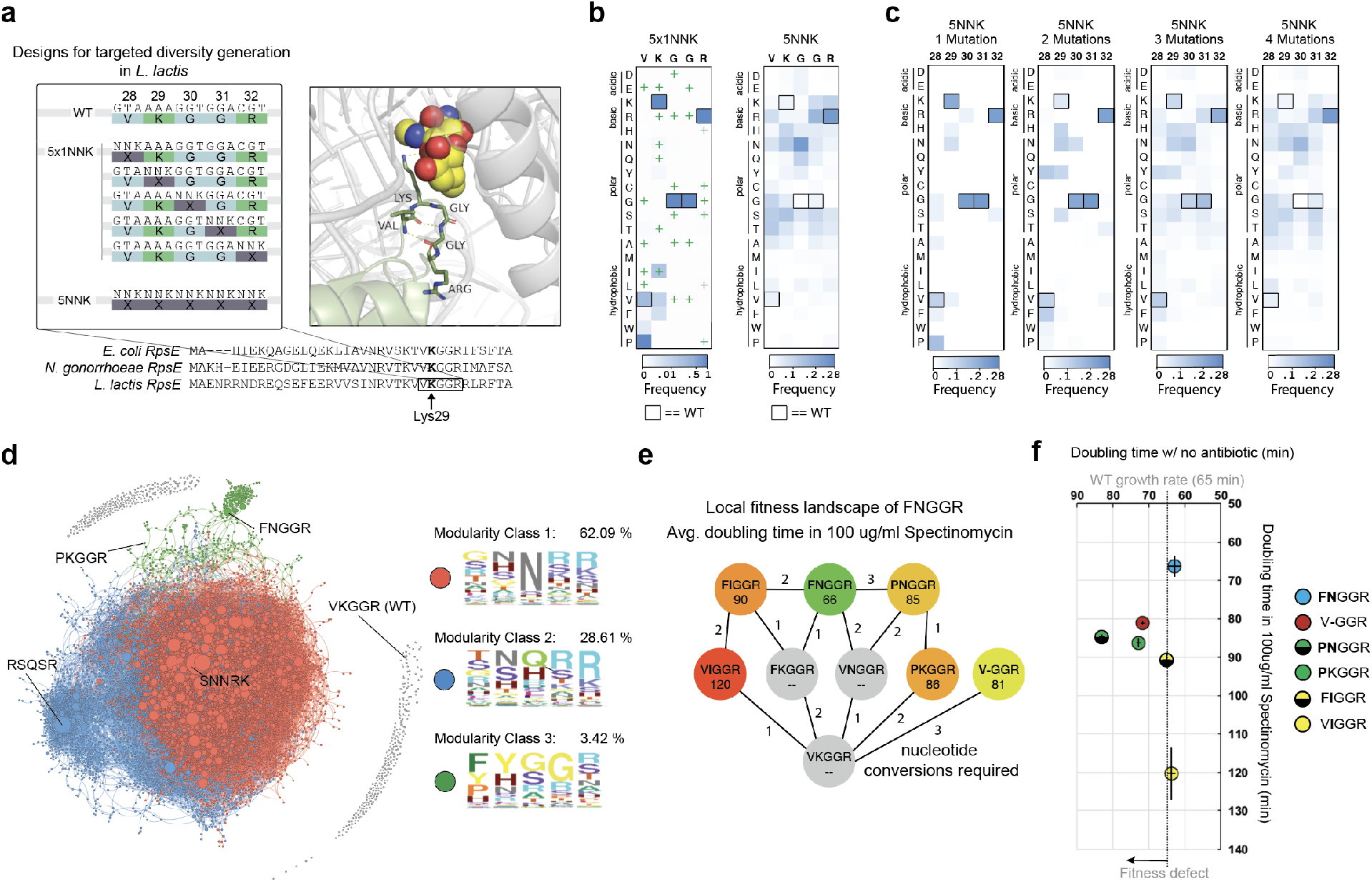
(a), Oligonucleotide design strategy. A sequence alignment of RpsE between *E. coli, N. gonorrhoeae* (a pathogen targeted by spectinomycin), and *L. lactis* shows a conserved 5AA region around the spectinomycin binding pocket (*E. coli* Lys26). 5 oligos can be pooled to introduce single degenerate codons at each amino acid position (5×1NNK), and a single oligo containing a fully degenerate sequence (5NNK) can be used to diversify the entire region. A crystal structure of *E. coli* RpsE bound to spectinomycin^29^ shows the approximate location of the antibiotic relative to the 5 targeted residues (Fig S15). (b), Normalized heat maps after selection and enrichment for the 5×1NNK single-amino acid saturation mutagenesis experiment vs. 5NNK combination mutagenesis experiment. WT residues are outlined in black. Amino acids accessible through a single-nucleotide conversion are indicated by a “+” (c), Heat maps of amino acid enrichment for single, double, triple and quadruple mutants found within the 5NNK mutant library. (d), Force directed graph of all spectinomycin resistant combination mutants with at least 10 reads, lines connect variants that could be accessed through a single-nucleotide mutation, and the size of dots reflects the relative enrichment (e), Shortest paths to highly enriched double mutant FNGGR, as well as the nucleotide conversions required (f), Fitness of selected mutants in the presence and absence of antibiotic. Error bars represent the standard deviation of four replicates.

After enrichment of resistant mutants by passaging, we used next-generation sequencing to identify the frequency of individual variants. Previously, error-prone mutagenesis in *E. coli* found one amino acid mutation (G30D in *L. lactis*) with lower levels of resistance, as well as a number of more resistant mutants formed through amino-acid deletions^17^. Our single-amino acid saturation mutagenesis library (5×1NNK) revealed ~15 new candidate resistant amino acid variants, with a great over-representation of a single mutant **P**KGGR, which requires two simultaneous nucleotide conversions (Fig 5b). In the combination mutant library, we found over 49,000 more candidate resistance mutants (Fig 5b). By generating heat maps of resistant variants containing 1, 2, 3 or 4 amino acid mutations relative to wild-type, we see that the enriched alleles change at different mutational depths (Fig 5c), indicating the presence of extensive epistatic effects. (When diversifying all 5 positions, the possibility of sampling WT amino acids allow us to identify single, double, triple, or quintuple mutants).

To analyze the combination mutant library holistically we generated a force-directed graph containing all mutants with at least 10 unique reads (Fig 5d). We connected each mutant to every other mutant that was accessible through a single-nucleotide mutation. Interestingly, none of these enriched mutants were accessible using single-nucleotide mutations from WT (Fig 5d). Most of the resistant variants clustered into two closely related groups, in which all 5 amino acids are enriched with polar or charged resides. Interestingly for a third cluster, WT residues are enriched at 3 of the 5 residues (“XX”GGR) (Fig 5d). We analyzed the local fitness landscape of FNGGR, the most enriched mutant in this cluster, and found that it was formed by the epistatic combination of two mutations with no individual effect (Fig 5e). Furthermore, we found that FNGGR, which would not be readily sampled through single nucleotide mutations, was more resistant than either the most resistant single mutant or a previously identified deletion mutant (Fig 5e). Additionally, although the most resistant single mutants and deletion mutants exhibited fitness defects, FNGGR grew with no appreciable fitness defect (Fig 5f). Due to the extensive epistasis at this locus, only methods like recombineering that traverse over local minima are able to properly characterize the genetic landscape and identify variants with extreme phenotypes. In this experiment, the high-editing efficiencies achievable via recombineering in *L. lactis* allowed us to simultaneously diversify 5 amino acid positions. In species with ~1000 (32^2) fold lower oligonucleotide incorporation efficiencies, similar experiments could be performed to diversify three neighboring amino-acid positions simultaneously.

## Discussion

In this work we examined the host-tropism of phage RecT proteins, which are used by microbiologists, bioengineers, and synthetic biologists for genome engineering, and which also —due to their high conservation across dsDNA phages^31^ — are likely key players in phage recombination and replication pathways^23^. We found that an interaction between RecT proteins and the host SSB largely determines their genome editing activity when moved between disparate bacterial species. Using chimeric SSBs we found that this RecT-SSB interaction depends on recognition of the 7 C-terminal amino acids of SSB. Although the RecT-SSB interaction is specific, we found some RecT proteins have a naturally broad host range (PapRecT) and interact with multiple distinct SSB tails. We also found that co-expression of an exogenous bacterial SSB further broadens the host-range of RecT proteins, and in certain species enables RecT protein functionality even if there is no basal host compatibility. We used these insights to develop recombineering in *L. rhamnosus*, improving oligonucleotide incorporation frequency over 2,500-fold, and *C. crescentus* improving oligonucleotide incorporation 870-fold.

We then demonstrated how the capabilities of recombineering surpass more broadly used error-prone methods of genome diversification by identifying mutants of RpsE in *L. lactis* that conferred spectinomycin resistance. We performed saturation mutagenesis across 5 amino acids neighboring the antibiotic binding pocket, as well as multiplexed mutagenesis to generate nearly all combination mutants across the same 5 positions. The enriched variants would have been difficult to identify with even the most efficient nucleotide diversification methods due to extensive epistatic effects within the mutational landscape. The path to the fittest characterized double mutant, **FN**GGR, would require either 3 simultaneous nucleotide conversions in two neighboring amino acids, or a path of 1 single, and then two double nucleotide conversions (Fig 5e). Because degeneracy can be introduced in the oligonucleotides, a single oligo pool which cost less than $100 was used to perform multiplexed mutagenesis and produce over 49,000 new resistance genotypes. These methods will likely be useful for interrogating other protein-small molecule, or protein-protein interaction domains in gene’s native context, and identify new phenotypes not observed through random mutagenesis. Although the work here focused on a selectable phenotype, non-selectable phenotypes such as fitness could also be examined as long as the sequencing depth is scaled by the efficiency of editing. In *L. lactis* with a relatively high editing efficiency of ~2%, 50x additional sequencing coverage would be needed to examine the same space of non-selected mutants. With saturation mutagenesis, where single amino acids are mutated, non-selectable phenotypes could be analyzed with an efficiency 1000-fold lower than in *L. lactis* with the same sequencing coverage.

There are many questions which remain about the mechanism of recombineering. First, there is no crystal structure of any full length RecT protein, and a structural understanding of the steps involved in oligonucleotide binding and annealing in the presence of SSB should be characterized to better understand this mechanism. Second, while co-expression of an exogenous SSB can expand the host range of RecT proteins, this is not a fully generalizable method, which raises questions about what other components may be involved in the mechanism. Possible causes of RecT-SSB portability failure may include: 1) an inhibitory interaction between RecTs and the host SSB (there are a number of examples in Fig 3a, b where an SSB reduced the efficiency of an otherwise functional RecT protein), and 2) SSB-induced toxicity (neither PapRecT + PaSSB, or Redβ + PaSSB could be transformed and expressed in *L. rhamnosus*). With the long-term goal of reliably establishing high efficiency homologous recombination methods in all bacterial species, understanding the mechanisms prohibiting the portability of RecT-SSB pairs will further help to expand and generalize these methods.

Although we focused here on ssDNA recombineering, we believe that other recombineering technologies enabled by RecT proteins can be established once a functional RecT protein is identified. For example, dsDNA recombineering, which is used to integrate larger constructs including genes and resistance markers, usually requires the presence of a cognate phage exonuclease. These proteins are almost always found within the phage operon containing the RecT, and can be readily co-expressed to enable dsDNA recombineering^21^. Surprisingly, we found that PapRecT + PaSSB enabled dsDNA recombineering in *L. lactis* even without including a cognate phage exonuclease (Fig S8), suggesting that SSB co-expression alone might be sufficient for dsDNA recombineering. Many of the other recombineering technologies rely on the presence of proteins that have very broad host range, and can likely be directly ported into species where functional RecT proteins are identified. These include technologies combining recombineering with Cas9, which are used for (1) the scarless selection of genomic edits (recombineering + Cas9 selection^32^), (2) simple barcoding methods for creating genome-wide mutant libraries (CREATE^7^), and (3) synthetic genome assembly (REXER^14^). This also includes technologies combining recombineering with the site specific recombinase BxB1, which is used to rapidly make selectable gene knockouts, insertions or fusions using oligonucleotide-incorporated attP sites (ORBIT^33^). We believe that our work, in addition to describing the nature of the phage RecT-host interaction, will help to spur the development of new recombineering technologies, and expedite the portability of these methods into new bacterial species.

## Materials and Methods

### Bacterial Strains and Culturing Conditions

The *E. coli* strain used was derived from EcNR2 with some modifications (EcNR2.dnaG_Q576A.tolC_mut.mutS::cat_mut.dlambda::zeoR)^6^. *L. lactis* strain NZ9000 was provided as a kind gift from Jan Peter Van Pijkeren. *M. smegmatis* strain mc(2)155 was purchased from ATCC. The *C. crescentus* strain used was NA1000.

All chemicals were purchased from Sigma Aldrich, unless stated otherwise. *E. coli* and its derivatives were cultured in Lysogeny broth - Low sodium (Lb-L) (10 g/L tryptone, 5 g/L yeast extract (Difco), PH 7.5 with NaOH), in a roller drum at 34 °C. *L. lactis* was cultured in M17 broth (Difco, BD BioSciences) supplemented with 0.5% (w/v) D-glucose, static at 30 °C. *M. smegmatis* was cultured in Middlebrook 7H9 Broth (Difco, BD BioSciences) with AD Enrichment (10x stock: 50g/L BSA, 20g/L D-glucose, 8.5 g/L NaCl), supplemented with glycerol and Tween 80 to a final concentration of 0.2% (v/v) and 0.05% (v/v), respectively, in a roller drum at 37 °C. *C. crescentus* was cultured in peptone-yeast extract (PYE) broth (2 g/L peptone, 1 g/L yeast extract (Difco), .3 g/L MgSO4, 0.5mM 0.5M CaCl2), shaking at 30 °C.

Plating was done on petri dishes of LB agar for *E. coli*, M17 Agar (Difco, BD BioSciences) supplemented with 0.5% (w/v) D-glucose for *L. lactis*, 7H10 (Difco, BD BioSciences) supplemented with AD Enrichment and 0.2% (v/v) glycerol for *M. smegmatis*, and PYE agar for *C. crescentus*. Antibiotics were added to the media when appropriate, at the following concentrations: 50 μg/mL carbenicillin for *E. coli,* 10 μg/mL chloramphenicol for *L. lactis*, and 100 μg/mL hygromycin B for *M. smegmatis*, 5 μg/ml kanamycin for *C. crescentus*. For the selective plates used to determine allelic recombination frequency, antibiotics were added as follows: 0.005% SDS for *E. coli*, 50 μg/mL rifampicin for *L. lactis*, 20 μg/mL streptomycin for *M. smegmatis*, and 5 μg/ml rifampicin for *C. crescentus*.

### Construction and Transformation of Plasmids

Plasmids were constructed using PCR fragments and Gibson Assembly. All primers and genes were obtained from Integrated DNA Technologies (IDT). Plasmids were derived from pARC8 for use in *E. coli* ^34^, pjp005 for use in *L. lactis* - a gift from Jan Peter Van Pijkeren^20^, pKM444 for use in *M. smegmatis* - a gift from Kenan Murphy (Addgene plasmid # 108319), and pBXMCS-2 for use in *C. crescentus* ^35^. Genes were codon optimized for each of the host organisms using IDT’s online Codon Optimization Tool. *E. coli* and *L. lactis* plasmid constructs were Gibson assembled, then directly transformed into electrocompetent *E. coli* and *L. lactis* strains. *M. smegmatis* plasmids were first cloned in NEB 5-alpha Competent *E. coli* (New England Biolabs) for plasmid verification before transformation into electrocompetent *M. smegmatis*. All cloning was verified by Sanger sequencing (Genewiz). Plasmids will be deposited in Addgene. All data is available from the authors upon reasonable request.

### Protein purification

To prepare Redβ for *in vitro* analysis, it was first cloned by Gibson cloning into pET-53-DEST, with a 6x poly-histidine tag followed by a glycine-serine linker and a TEV protease site (MHHHHHHGSGENLYFQG) appended to its N-terminus. After purification and treatment with TEV protease, this leaves only an N-terminal glycine before the start codon. Overnight cultures of *E. coli* BL21 (DE3) (NEB) with the expression construct were diluted 1:100 into Fernbach flasks, grown to an OD of ~0.5, and induced with 1 mM IPTG at 37 °C for 4 h. Cultures were pelleted at 10,000 × g in a fixed angle rotor for 10 min and the supernatant decanted. Bacterial pellets were resuspended in 30 mL of lysis buffer (150 mM NaCl, 0.1% v/v Triton-X, 50 mM TRIS-HCl pH 8.0) and sonicated at 80% power, 50% duty cycle for 5 minutes on ice. The lysed cultures were again centrifuged for 10 min at 15,000 × g in a fixed angle rotor. The supernatant was then incubated for 30 minutes at room temperature with HisPur cobalt resin (Thermo) and column purified on disposable 25 ml polypropylene columns (Thermo). The protein-bound resin was washed with four column volumes of wash buffer (150 mM NaCl, 10 mM imidazole, 50 mM TRIS-HCl pH 8.0) and bound protein was eluted with two column volumes of elution buffer (150 mM NaCl, 250 mM imidazole, 50 mM TRIS-HCl pH 8.0). Protein eluates were dialyzed overnight against 25 mM TRIS-HCl pH 7.4 with 10,000 MWCO dialysis cassettes (Thermo), concentration was measured by Qubit (Thermo) and 1.5 mg of protein was cleaved in a 2 ml reaction with 240 Units of TEV protease (NEB) for two hours at 30 °C. The TEV cleavage reaction was re-purified with cobalt resin, except that in this case the flow-through was collected, as the His tag and the TEV protease were bound to the resin. Expression and successful TEV cleavage were confirmed by SDS-PAGE. Protein was concentrated in 10,000 MWCO Amicon protein concentrators (Sigma), protein concentration was assayed by Qubit, and an equal volume of glycerol was added to allow storage at −20 °C. *E. coli* and *L. lactis* SSBs were prepared according to previously published protocol (Lohman, Green, and Beyer, 1986) without the use of an affinity tag^36^.

### Oligonucleotide annealing and quenching experiments

Fluorescent (tolC-r.null.mut-3’FAM) and quenching (tolC-f.null.mut-5’IBFQ) oligos were ordered from Integrated DNA Technologies. Unless otherwise indicated, 50 nM of each oligo was incubated in 25 mM TRIS-HCl pH 7.4 with 1.0 μM *Ec*_SSB or *Ll*_SSB at 30 °C for 30 minutes. 100 μl of each oligo mixture were then combined into a 96-well clear-bottom black assay plate (Costar), incubated a further 60 minutes at 30 °C, and annealing was tracked on a Synergy H4 microplate reader (BioTek) with fluorescence excitation set to 495 nm and emission set to 520 nm. After 60 minutes, 20 μl of a solution with or without 25μM Redβ and containing 100 mM MgCl_2_ was added to achieve a final reaction concentration of 2.5 μM Redβ and 10 mM MgCl_2_. The annealing was then tracked over 10 hours in a the Synergy H4 microplate reader with the setting indicated above.

### Preparation of Electrocompetent *E. coli*

A single colony of *E. coli* was grown overnight to saturation. In the morning 30 μL of dense culture was inoculated into 3 mL of fresh media and grown for 1 hour. To induce gene expression of the pARC8 vector for recombineering experiments, L-arabinose was added to a final concentration of 0.2% (w/v) and the cells were grown an additional hour. 1 mL of cells were pelleted at 4 °C by centrifugation at 12,000 x g for 2.5 minutes and washed twice with 1 mL of ice-cold dH_2_O. Cells were resuspended in 50 μL ice-cold dH_2_O containing DNA and transferred to a pre-chilled 0.1 cm electroporation cuvette.

### Preparation of Electrocompetent *L. lactis*

A single colony of *L. lactis* was grown overnight to saturation. 500 μL of dense culture was inoculated into 5 mL of fresh media, supplemented with 500mM sucrose and 2.5% (w/v) glycine, and grown for 3 hours. To induce gene expression of the pJP005 vector for recombineering experiments, the cells were grown for an additional 30 min after adding 1 ng/mL freshly diluted nisin, unless stated otherwise. For the optimized condition (Fig 4b), 10 ng/mL nisin was used. Cells were pelleted at 4 °C by centrifugation at 5,000 x g for 5 minutes and washed twice with 2 mL of ice-cold electroporation buffer (500mM sucrose containing 10% (w/v) glycerol) by centrifugation at 13,200 x g for 2.5 minutes. Cells were resuspended in 80 μL ice-cold electroporation buffer containing DNA and transferred to a pre-chilled 0.1 cm electroporation cuvette.

### Preparation of Electrocompetent *M. smegmatis*

A single colony of *M. smegmatis* was grown overnight to saturation. The next day 25 μL of dense culture was inoculated into 5 mL of fresh media in the evening and grown overnight to an OD_600_of 0.9. Cells were pelleted at 4 °C by centrifugation at 3,500 x g for 10 minutes and washed twice with 10 mL ice-cold 10% glycerol. Cells were resuspended in 360 μL ice-cold 10% glycerol and transferred along with 10 μL of DNA to a pre-chilled 0.2 cm electroporation cuvette.

### Preparation of Electrocompetent *C. crescentus*

A single colony of *C. crescentus* was grown overnight. The next day cells were diluted back to OD ~0.001 in 25 mL PYE, and grown overnight. The next day, 250 μL of 30% xylose was added to cells at OD ~0.2. Cells were harvested at between OD = 0.5 and OD = 0.7, spun at 10,000 rpm for 10 min, and then washed twice in 12.5ml of ice-cold dH_2_O, washed once in 12.5ml of ice-cold 10% glycerol, then washed and resuspended in 2.5ml of ice-cold 10% glycerol. 90 μL of cells were added along with DNA to 0.1cm cuvettes and incubated on ice for 10 min.

### Recombineering Experiments

Electrocompetent cells were electroporated with 90-mer oligos at: 1 uM for *E. coli*, 50 μg for *L. lactis*, and 10uM for *C. crescentus*. 70-mer oligos were used at 1 μg for *M. smegmatis*. All oligos were obtained from IDT and can be found under “Oligonucleotides for genome editing” in materials and methods. For dsDNA experiments *L. lactis* was electroporated with 1.5 μg purified linear dsDNA. Cells were electroporated using a Bio-Rad gene pulser set to 25 μF, 200 Ω, and 1.8 kV for *E. coli*, 2.0 kV for *L. lactis*, and 1.5kV for *C. crescentus* and to 1000 Ω and 2.5 kV for *M. smegmatis*. Immediately after electroporation, cells were recovered in fresh media for 3 hours for *E. coli*, 1 hour for *L. lactis*, overnight for *M. smegmatis* and overnight for *C. crescentus*. *L. lactis* recovery media was supplemented with MgCl_2_and CaCl_2_ at a concentration of 20 mM and 2 mM, respectively. *E. coli* recovery media was supplemented with carbenicillin. *M. smegmatis* recovery media was supplemented with hygromycin. *C. crescentus* recovery media was supplemented with 0.3% xylose and kanamycin. After recovery, the cells were serial diluted and plated on non-selective vs. selective agar plates to obtain approximately 50-500 CFU/plate. Colonies were counted using a custom script in Fiji, and allelic recombination frequency was calculated by dividing the number of colonies on selective plates, with the number of colonies on non-selective plates.

### Next-generation sequencing and analysis for *L. lactis* Spectinomycin resistant mutants

For the site-saturation experiment (1×5NNK) half of the cells recovered from a single transformation were used. For the combinatorial experiment (5NNK) cells recovered from 30 parallel transformations were used. After transformation, cells were recovered for 1hr, then plated on 100 μg/mL Spectinomycin GM17 plates. After 2 days of growth, cells were resuspended in 1 mL of culture per plate, and recovered overnight. The cultures were then passaged to enrich for resistant mutants. During passaging for 1×5NNK: 100 μL confluent cells were passaged into 10 mL culture and grown overnight for the single-amino acid library, and for 5NNK: 500 μL confluent cells were passaged into 50 mL culture and grown overnight for the combinatorial mutant library. Cultures were used as templates for initial amplification using the *L. lactis* RpsE locus primers given under “Next-generation sequencing of spectinomycin resistance mutants (*L. lactis*)”. Cultures were collected at various time points: 1 hr after electroporation and after overnight recovery in Spectinomycin for the 1NNK library, and 1 hr after electroporation as well as after 1, 2, and 3 overnight passages in Spectinomycin for the 5NNK library. The data from 3 overnight passages was used for subsequent analysis. 50 μL of cells at each timepoint were spun down to make a pellet and used in a 50 μL PCR reaction. Initial amplification was monitored by qPCR for ~20 cycles until late exponential phase. A second round of qPCR was used to add indexing primers for Illumina sequencing, and run for ~15 cycles until late exponential phase. Samples were quantified by Qubit, combined and sequenced using Illumina MiSeq with reagent kit V3. Sequencing was done using a single 150bp read using RpsE locus primer R. For each timepoint, sequencing reads were processed using a python script, filtered such that reads matched the genomic locus 21 bp upstream and 21 bp downstream of the target site.

### *E. coli* RecT-SSB Toxicity Assay

A single colony of each construct for *E. coli* was grown overnight in a 96-well plate containing 150 μL of media per well with carbenicillin. 1.5 μL overnight culture was inoculated into 150 μL fresh media and grown for 2 hours to reach exponential phage. 1.5 μL of exponential culture was re-inoculated into 150 μL fresh media containing .2% (v/v) L-arabinose to induce expression of the genes. OD600 was then measured every three minutes, for a time course of 7 hours in a BioTek Eon plate reader, set to 34 °C with double orbital shaking. Growth curves were analyzed in Matlab using a custom script to determine the doubling time.

### *L. lactis* Spectinomycin and wildtype fitness assay

To assay spectinomycin resistance in *L. lactis*, single colonies were grown in 150 μL overnight with no antibiotic in 96 well plates. In the morning cells 1.5 μL confluent culture was then inoculated into 300 μL media containing no antibiotic or 100 μg/mL Spectinomycin in 96 well black assay plates with clear bottoms. These plates were covered with a “Breathe-Easy” sealing film, and cells were grown at 30C for 7 hours in a BioTek Eon plate reader, with OD600 measured every 3 minutes.

### Generating the force-directed graph of spectinomycin resistance mutants

Using python, an edges matrix was generated between SpecR combination mutants that had more than 10 reads and were accessible through single nucleotide mutations. This matrix was uploaded to the network visualization software Gephi, and a separate node matrix of the SpecR mutants was uploaded to add the genotype labels and enrichment values. Using Gephi, a force directed graph was produced using Force Atlas 2 (scaling = 25, gravity = 250). Once the nodes reached steady-state locations, the modularity statistic was performed using a resolution of 3.0 to identify the major clusters.

### Statistics

All error bars represent standard deviation. Statistical tests are given in table S1.

### Code Availability

The custom scripts used to count colonies in Fiji, analyze doubling times in MATLAB, analyze Illumina sequencing data, and generate the adjacency matrix for the force-directed graph are available upon request.

### Protein structures

Protein structure images (Fig 2a) were downloaded from PyMOL: Schrodinger LLC, The PyMOL Molecular Graphics System, Version 1.8 (2015).

## Acknowledgments

We thank Gleb Kuznetsov and Georgia Squyres for reviewing the manuscript and providing helpful feedback. **Funding:** This work was supported through National Institute of General Medical Sciences under Grant No. 1U01GM110714-01 and the Department of Energy DE-FG02-02ER63445 to G.M.C.. This work was also supported by NIH grant R01GM082899 to M.T.L. who is an Investigator of the Howard Hughes Medical Institute. G.T.F. was supported by the National Science Foundation Graduate Research Fellowship under Grant No. DGE1745303. D.A.S was supported by a Landry Cancer Biology Research Fellowship. K.G. was supported by the National Science Foundation Graduate Research Fellowship under Grant No. DGE1745302. **Author contributions:** G.T.F., T.M.W., and G.M.C. conceived the study; G.T.F. designed the bacterial experiments; T.M.W., F.B.P., I.D.L., and J.Z assisted with bacterial experimental design; G.T.F., F.B.P., I.D.L., and J.Z. carried out the experiments in *E. coli, L. lactis,* and *M. smegmatis* and analyzed the data; D.A.S. performed bioinformatic analysis; K.G. and G.T.F performed *C. crescentus* experiments under the supervision of M.T.L and G.M.C; T.M.W., designed and performed the in-vitro experiments; A.D. contributed to experiments in *L. rhamnosus;* H.K., V.V. and S.W. assisted with experiments; G.T.F. wrote the manuscript with input from all other authors; G.T.F and F.B.P. generated the figures; X.R., C.J.G, and M.J.L, contributed to project conception, S.L.S and J.A. provided supervision; G.M.C. supervised the study. **Competing interests:** G.M.C. is a founder of 64-x, Enevolv, and GRO Biosciences. G.T.F, T.M.W., and G.M.C. are named inventors on a patent application related to the technologies described in this article. Other potentially relevant financial interests are listed at http://arep.med.harvard.edu/gmc/tech.html. **Data and materials availability:** Nextgeneration sequencing data will be deposited into the National Center for Biotechnology Information Sequence Read Archive database. Plasmids will be available under a materials transfer agreement from Addgene.

