## Supplementary Data for "Characterizing the portability of RecT-mediated oligonucleotide recombination"

**This PDF file includes:**

Oligonucleotide and Gene sequences
Supplementary Note S1 to S3
Figures S1 to S15
Supplementary Table S1

Oligonucleotides for genome editing

- *Lactococcus Lactis*

| | |
|---|---|
| Rifampicin resistance 4bp mismatch: | C*G*AgagataccaccaggtcctaaggcagagaaacgacgtttgtTGCTaagctcagacaaaggattatgttggtccataaattgtgacaac |
| Rifampicin resistance 1bp mismatch: | C*G*AgagataccaccaggtcctaaggcagagaaacgacgtttgtTtgaaagctcagacaaaggattatgttggtccataaattgtgacaac |
| Streptomycin resistance (K56R): | G*A*AAGACGTACACGCGCGAATTTACGAAGCGCTGAGTTAGGTTTTCTAGGAGTCATTGTACCAACACGAGTTGCTACTCCACGTTTTTGT |
| NNKx5 oligo Spectinomycin resistance | C*G*GTCACCAACAACAACAAGTGCTGCAAAGCGAAGACGMNNMNNMNNMNNMNNAACTTTAGTAACGCGGTTAATTGAAACTACGCGTTCT |
| 5x1NNK oligo Spectinomycin resistance (AA 32) | C*G*GTCACCAACAACAACAAGTGCTGCAAAGCGAAGACGMNNTCCACCTTTTACAACTTTAGTAACGCGGTTAATTGAAACTACGCGTTCT |
| 5x1NNK oligo Spectinomycin resistance (AA 31) | C*G*GTCACCAACAACAACAAGTGCTGCAAAGCGAAGACGACGMNNACCTTTTACAACTTTAGTAACGCGGTTAATTGAAACTACGCGTTCT |
| 5x1NNK oligo Spectinomycin resistance (AA 30) | C*G*GTCACCAACAACAACAAGTGCTGCAAAGCGAAGACGACGTCCMNNTTTTACAACTTTAGTAACGCGGTTAATTGAAACTACGCGTTCT |
| 5x1NNK oligo Spectinomycin resistance (AA 29) | C*G*GTCACCAACAACAACAAGTGCTGCAAAGCGAAGACGACGTCCACCMNNTACAACTTTAGTAACGCGGTTAATTGAAACTACGCGTTCT |
| 5x1NNK oligo Spectinomycin resistance (AA 28) | C*G*GTCACCAACAACAACAAGTGCTGCAAAGCGAAGACGACGTCCACCTTTMNNAACTTTAGTAACGCGGTTAATTGAAACTACGCGTTCT |
| Spectinomycin resistance mutation matching *E. coli* (G27D matches G30D) | C*G*GTCACCAACAACAACAAGTGCTGCAAAGCGAAGACGACGTCCATCTTTTACAACTTTAGTAACGCGGTTAATTGAAACTACGCGTTCT |
| NNKx5 oligo Spectinomycin resistance Validation 1 (RTNAR) | C*G*GTCACCAACAACAACAAGTGCTGCAAAGCGAAGACG CCGCGCATTCGTACG AACTTTAGTAACGCGGTTAATTGAAACTACGCGTTCT |
| NNKx5 oligo Spectinomycin resistance Validation 2 (NGTRF) | C*G*GTCACCAACAACAACAAGTGCTGCAAAGCGAAGACG AAACCTAGTACCATT AACTTTAGTAACGCGGTTAATTGAAACTACGCGTTCT |
| 5x1NNK oligo Spectinomycin resistance Validation 1 (V28P) | C*G*GTCACCAACAACAACAAGTGCTGCAAAGCGAAGACGACGTCCACCTTTAGGAACTTTAGTAACGCGGTTAATTGAAACTACGCGTTCT |
| 5x1NNK oligo Spectinomycin resistance Validation 2 (K29I) | C*G*GTCACCAACAACAACAAGTGCTGCAAAGCGAAGACGACGTCCACCAATTACAACTTTAGTAACGCGGTTAATTGAAACTACGCGTTCT |
| Spectinomycin resistance validation R at 28 | C*G*GTCACCAACAACAACAAGTGCTGCAAAGCGAAGACGACGTCCACCTTTCCTAACTTTAGTAACGCGGTTAATTGAAACTACGCGTTCT |

| | |
|---|---|
| Spectinomycin resistance validation R at 28, NN at 30-31 | C*G*GTCACCAACAACAACAAGTGCTGCAAAGCGAAGACGACGATTATTTTTCCTAACTTTAGTAACGCGGTTAATTGAAACTACGCGTTCT |
| Spectinomycin resistance validation R at 28, N at 31 | C*G*GTCACCAACAACAACAAGTGCTGCAAAGCGAAGACGACGATTACCTTTCCTAACTTTAGTAACGCGGTTAATTGAAACTACGCGTTCT |
| Spectinomycin resistance validation R at 28, N at 30 | C*G*GTCACCAACAACAACAAGTGCTGCAAAGCGAAGACGACGTCCATTTTTCCTAACTTTAGTAACGCGGTTAATTGAAACTACGCGTTCT |
| Spectinomycin resistance validation NN at 30-31 | C*G*GTCACCAACAACAACAAGTGCTGCAAAGCGAAGACGACGATTATTTTTACAACTTTAGTAACGCGGTTAATTGAAACTACGCGTTCT |
| Spectinomycin resistance validation N at 31 | C*G*GTCACCAACAACAACAAGTGCTGCAAAGCGAAGACGACGATTACCTTTTACAACTTTAGTAACGCGGTTAATTGAAACTACGCGTTCT |
| Spectinomycin resistance validation N at 30 | C*G*GTCACCAACAACAACAAGTGCTGCAAAGCGAAGACGACGTCCATTTTTACAACTTTAGTAACGCGGTTAATTGAAACTACGCGTTCT |
| Spectinomycin resistance validation R at 31 | C*G*GTCACCAACAACAACAAGTGCTGCAAAGCGAAGACGACGCCTACCTTTTACAACTTTAGTAACGCGGTTAATTGAAACTACGCGTTCT |
| Spectinomycin resistance validation N at 30, R at 31 | C*G*GTCACCAACAACAACAAGTGCTGCAAAGCGAAGACGACGCCTATTTTTACAACTTTAGTAACGCGGTTAATTGAAACTACGCGTTCT |
| Spectinomycin resistance validation MNR at 29-31 | C*G*GTCACCAACAACAACAAGTGCTGCAAAGCGAAGACGACGCCTATTCATTACAACTTTAGTAACGCGGTTAATTGAAACTACGCGTTCT |
| Spectinomycin resistance validation M at 29 | C*G*GTCACCAACAACAACAAGTGCTGCAAAGCGAAGACGACGTCCACCCATTACAACTTTAGTAACGCGGTTAATTGAAACTACGCGTTCT |
| Spectinomycin resistance validation M at 29, R at 31 | C*G*GTCACCAACAACAACAAGTGCTGCAAAGCGAAGACGACGCCTACCCATTACAACTTTAGTAACGCGGTTAATTGAAACTACGCGTTCT |
| Spectinomycin resistance validation M at 29, N at 30 | C*G*GTCACCAACAACAACAAGTGCTGCAAAGCGAAGACGACGTCCATTCATTACAACTTTAGTAACGCGGTTAATTGAAACTACGCGTTCT |
| Spectinomycin resistance validation SNNRK | C*G*GTCACCAACAACAACAAGTGCTGCAAAGCGAAGACGTTTACGATTATTTGAAACTTTAGTAACGCGGTTAATTGAAACTACGCGTTCT |
| Spectinomycin resistance validation FNGGR | C*G*GTCACCAACAACAACAAGTGCTGCAAAGCGAAGACGACGTCCACCGTTAAAAACTTTAGTAACGCGGTTAATTGAAACTACGCGTTCT |
| Spectinomycin resistance validation FKGGR | C*G*GTCACCAACAACAACAAGTGCTGCAAAGCGAAGACGACGTCCACCTTTAAAAACTTTAGTAACGCGGTTAATTGAAACTACGCGTTCT |

| | |
|---|---|
| Spectinomycin resistance validation VNGGR | C*G*GTCACCAACAACAACAAGTGCTGCAAAGCGAAGACGACGTCCACCGTTTACAACTTTAGTAACGCGGTTAATTGAAACTACGCGTTCT |
| Spectinomycin resistance validation PNGGR | C*G*GTCACCAACAACAACAAGTGCTGCAAAGCGAAGACGACGTCCACCGTTCGGAACTTTAGTAACGCGGTTAATTGAAACTACGCGTTCT |
| Spectinomycin resistance validation V-GGR | C*G*GTCACCAACAACAACAAGTGCTGCAAAGCGAAGACGACGTCCACCTACAACTTTAGTAACGCGGTTAATTGAAACTACGCGTTCT |
| Spectinomycin resistance validation FIGGR | C*G*GTCACCAACAACAACAAGTGCTGCAAAGCGAAGACGACGTCCACCAATAAAAACTTTAGTAACGCGGTTAATTGAAACTACGCGTTCT |

- *Escherichia coli*

| | |
|---|---|
| 4bp mutation conferring SDS resistance (tolC targeting) | A*G*CAAGCACGCCTTAGTAACCCGGAATTGCGTAAGTCTGCCGCTAAATCGTGATGCTGCCTTTGAAAAATTAATGAAGCGCGCAGTCCA |

- *Mycobacterium smegmatis*

| | |
|---|---|
| 1bp Streptomycin resistance (targets rpsL): | G*T*CAGCTTCACGCGCGCGACCTTCCGGAGCGCCGAGTTCGGCTTCCTCGGAGTGGTGGTGTAAACGCGCGTGCACACGCCGCGACGCTGC |

- *Lactobacillus rhamnosus*

| | |
|---|---|
| 3bp rifampacin resistance (targets RpoB): | A*C*GAGTCAAGCCACCAGGTCCAAGGGCAGACAGACGCCGTTTGTCCGTTAATTCGCCCAACGGATTGGTCTGATCATGAACTGTGACAG |

- *Caulobacter crescentus*

| | |
|---|---|
| Rifampicin resistance 4bp mismatch: | C*G*CGGGTCAGACCGCCCGGGCCGAGGGCCGAGAGACGACGCTTCCTCGTGATTTCCGACAGCGGGTTCGTCTGGTCCATGAACTGCGACA |
| Rifampicin resistance 1bp mismatch: | C*G*CGGGTCAGACCGCCCGGGCCGAGGGCCGAGAGACGACGCTTGCGGGTGATTTCCGACAGCGGGTTCGTCTGGTCCATGAACTGCGACA |

Oligos used for fluorescence quenching:

| | |
|---|---|
| Quenching oligo: tolC-f.null.mut-5'IBFQ | /5’IBFQ/TGGACTGCGCGCTTCATTAATTTTTCAAAGGCAGCATCACGATTTAGCGGCAGACTTACGCAATTCCGGGTTACTAAGGCGTGCTTGCT |
| Fluorescent oligo: tolC-r.null.mut-3'6FAM | AGCAAGCACGCCTTAGTAACCCGGAATTGCGTAAGTCTGCCGCTAAATCGTGATGCTGCCTTTGAAAAAATTAATGAAGCGCGCAGTCCA/3’FAM/ |

Primers

- MASC PCR (*L. lactis*)

| | |
|---|---|
| Rifampacin 1bp PCR F WT | AGAGAAACGACGTTTGTG |
| Rifampacin 1bp PCR F Mut | CAGAGAAACGACGTTTGTT |
| Rifampacin 1bp PCR R Common | CGTTCAGTTGGTGAATTACT |

- dsDNA validation of genome integration (*L. lactis*)

| | |
|---|---|
| Colony PCR primer for integration into genome F | GTAGATAAATTATTAGGTATACTACTGACAGC |
| Colony PCR primer for integration into genome R | CGTACCAAGCTCAGTTAATTG |

- Sanger sequencing for spectinomycin resistance mutants (*L. lactis*)

| | |
|---|---|
| L. lactis RpsE sequencing primer R | ACACTTGCATCAGCTTCATC |
| L. lactis RpsE sequencing primer F | ATTTTGATGTTACGTCAGCAACAC |

- Next-generation sequencing of spectinomycin resistance mutants (*L. lactis*)

| | |
|---|---|
| Next gen. seq. primer Spectinomycin resistance F 1 | ctttccctacacgacgctcttccgatctATGGCAGAAAACAGAAGAAATG |
| Next gen. seq. primer Spectinomycin resistance F 2 | ctttccctacacgacgctcttccgatctNATGGCAGAAAACAGAAGAAATG |
| Next gen. seq. primer Spectinomycin resistance F 3 | ctttccctacacgacgctcttccgatctNNATGGCAGAAAACAGAAGAAATG |
| Next gen. seq. primer Spectinomycin resistance F 4 | ctttccctacacgacgctcttccgatctNNNATGGCAGAAAACAGAAGAAATG |
| Next gen. seq. primer Spectinomycin resistance R | ggagttcagacgtgtgctcttccgatctTTTTACCACCACCGAAGAC |

Nucleotide sequences of codon optimized Genes: RecT

- *L. lactis* (The same genes are used for *L. rhamnosus*)

| | |
|---|---|
| λ Beta | ATGAGTACTGCACTTGCAACATTAGCTGGCAAGTTAGCAGAGCGTGTTGGTATGGATTCAGTCGACCCTCAGGAGCTTATAACTACCTTACGTCAAACAGCGTTCAAGTGTGACGCCTCTGATGCACAATTTATCGCTTTGCTTATCGTAGCTAACCAGTATGGGTTGAATCCTTGGACGAAGGAGATATACGCTTTCCCGGATAAGCAGAACGGTATTGTTCCTGTAGTAGGTGTCGATGGATGGAGTAGAATTATCAATGAAAATCAACAGTTCGATGGCATGGACTTCGAGCAGGATAATGAATCATGTACCTGCCGTATATATAGAAAAGACCGAAATCACCCAATTTGTGTGACTGAATGGATGGATGA |

| | |
|---|---|
| | GTGCAGACGTGAGCCGTTCAAGACCCGAGAAGGCCGTGAAATCACT GGTCCGTGGCAATCACATCCAAAGAGAATGTTGCGTCACAAGGCGA TGATTCAGTGCGCCCGTTTAGCTTTTGGGTTTGCTGGCATTTACGACA AGGACGAAGCTGAAAGAATCGTTGAAAACACTGCATATACCGCTGA ACGACAACCGGAGCGTGACATTACGCCAGTGAATGACGAGACAATG CAGGAAATTAACACGTTGTTGATTGCTTTGGACAAAACGTGGGACGA CGACTTGTTACCACTTTGTAGCCAAATTTTTCGTCGAGACATTAGAG CTTCATCTGAGCTTACACAAGCTGAAGCCGTCAAGGCATTGGGGTTT TTGAAACAAAAAGCTACCGAACAGAAGGTAGCGGCATAA |
| PapRecT | ATGGGAACCGCCCTTACACCTCTTTTGACAAAGTTCGCAACCAGATA TGAGATGGGAACGACCCCTGAAGAGGTTGCTAATACATTGAAACAA ACATGCTTCAAGGGACAGGTCAACGACAGTCAAATGGTAGCCCTTTT GATAGTCGCTGACCAGTACAAGTTAAACCCGTTCACCAAGGAGTTGT ATGCATTCCCTGACAAGAATAATGGAATCGTGCCAGTTGTTGGTGTC GATGGATGGGCGAGAATAATAAACGAGAACCCTCAGTTTGATGGTA TGGAATTTTCTATGGACCAGCAGGGCACTGAGTGCACGTGCAAAATC TATCGTAAGGATCGTTCTCACGCAATAAGCGCTACGGAATATATGGC CGAATGTAAGAGAAATACGCAACCTTGGCAAAGTCACCCACGACGT ATGTTAAGACATAAAGCCATGATCCAGTGTGCGCGATTAGCATTCGG CTTCGCTGGTATCTACGATCAAGACGAAGCGGAACGAATAGTCGAA AGAGACGTTACTCCGGCGGAGCAGTACGAAGATGTCAGCGAAGCTA TATGCTTGATTAAGGACAGCCCGACGATGGAGGATTTACAGGCAGC GTTCAGCAATGCCTGGAAGGCGTATAAAACCAAAGGTGCAAGAGAC CAATTGACAGCCGCCAAGGATCAGCGTAAAAAGGAATTACTTGATG CCCCAATAGATGTCGAGTTCGAAGAAACTGGCGATGATAGAGCAGC ATAA |
| MspRecT | ATGGCAGAAAACGCCGTGACGAAACAAGACTCACCTAAAGCCCCAG AAACGATATCACAGGTCCTTCAAGTGTTAGTACCTCAATTAGCTCGA GCCGTACCTAAGGGAATGGATCCTGATAGAATAGCTCGAATCGTCCA GACCGAGATCAGAAAGAGTAGAAATGCGAAAGCGGCGGGAATCGC CAAACAGTCATTGGATGACTGCACGCAAGAAAGCTTCGCCGGGGCG TTACTTACAAGTGCGGCATTGGGCCTTGAGCCAGGCGTTAACGGTGA GTGTTATCTTGTTCCATACAGAGACACAAGAAGAGGTGTCGTCGAGT GCCAGTTAATTATCGGGTATCAAGGAATCGTTAAATTGTTTTGGCAA CATCCGCGAGCCTCTCGAATAGACGCCCAGTGGGTGGGGGCAAACG ATGAATTCCATTATACAATGGGTCTTAACCCAACCTTAAAACATGTA AAGGCTAAGGGAGACCGAGGAAACCCAGTATATTTTTATGCTATCGT AGAGGTCACGGGTGCCGAGCCTTTGTGGGATGTCTTCACAGCTGATG AGATTAGAGAGTTGAGAAGAGGTAAAGTCGGTTCAAGCGGGGATAT TAAGGATCCGCAGAGATGGATGGAGCGAAAGACAGCGTTAAAACAA GTGCTTAAGCTTGCTCCAAAAACTACTCGTTTGGACGCAGCAATACG AGCGGACGATAGACCGGGAACAGATTTGTCTCAAAGCCAGGCGTTG GCATTACCTAGTACAGTTAAGCCAACAGCAGACTACATAGACGGTG AGATTGCAGAGCCACACGAGGTTGACACTCCGCCTAAAAGCAGTCG AGCACAACGAGCTCAGAGAGCGACTGCCCCAGCCCCAGACGTTCAG ATGGCCAATCCGGATCAATTAAAGCGTTTGGGAGAGATTCAAAAAG |

| | |
|---|---|
| | CCGAAAAGTACAATGATGCCGACTGGTTTAAGTTCTTAGCTGATAGT GCAGGGGTGAAAGCGACAAGAGCAGCTGATCTTACATTTGATGAAG CAAAAGCTGTAATAGATATGTTCGACGGCCCAAATGCTTGA |
| LrpRecT | ATGGCTAATCAAGTAGCACAACAGCAGAAACCGACTAAGCTAACCG ATCTTGTATTAGATCGTGTTAAACAAATGCAAGACACGCAGGACTTG TCACTACCCAAGAATTACAACGCTTCTAATGCGTTGAATGCAGCCTT TCTCGAATTACAAAAAGTACAAGACCGTAATCATCGGCCAGCCTTAG AAGTATGTTCTCATGACTCGATTGTTAAGTCCTTGTTAGATATGACAC TGCAAGGGCTATCCCCAGCAAAAGATCAATGCTACTTCATCGTATAC GGCAATGAGCTTCAAATGCAACGGAGCTATTTCGGTACTGTTGCAGC AGTTAAGCGACTGGATGGTGTTAAGAAAGTTAGGGCAGAAGTTGTT CACGAAAAAGATGACTTTGAAATTGGTGCTAATGAAGACATGGAGC TAGTCGTTAAGAGGTTCGTTCCTAAGTTTGAAAATCAAGATAATCAA ATTATTGGAGCTTTTGCCATGATTAAGACTGATGAAGGTACTGACTT TACTGTTATGACTAAGAAAGAGATTGATCAGTCATGGGCACAAACA CGTCAAAAAAATAACAAAGTACAGCAGAATTTTAGCCAAGAAATGG CAAAGCGTACTGTGCTTAATCGTGCCGCTAAGATGTTTATTAACACG TCTGATGATAGTGACCTTTTAACTGGTGCTATCAACGATACAACAAG CAACGAATACGATGATGAGCGTCGAGATGTAACGCCCGTTGAGGAT GAAAAACAAAGTACTGATAAATTGCTAGAAGGATTTCAAAAGTCAC AAGAAGCGAAGGCTAAGTGGGTAAGTAATGATGGCAACAGCAACGA AGGCAAAGAAACCAGTGAAGAAGTCGCAGACGGACAAACAGAACT CTTCAGCGAAGGGACAATCAAACCAGCCGATGAAGCTGACAGCTAA |

- *E. coli*

| | |
|---|---|
| λ Beta | ATGAGTACTGCACTCGCAACGCTGGCTGGGAAGCTGGCTGAACGTGT CGGCATGGATTCTGTCGACCCACAGGAACTGATCACCACTCTTCGCC AGACGGCATTTAAAGGTGATGCCAGCGATGCGCAGTTCATCGCATTA CTGATCGTTGCCAACCAGTACGGCCTTAATCCGTGGACGAAAGAAAT TTACGCCTTTCCTGATAAGCAGAATGGCATCGTTCCGGTGGTGGGCG TTGATGGCTGGTCCCGCATCATCAATGAAAACCAGCAGTTTGATGGC ATGGACTTTGAGCAGGACAATGAATCCTGTACATGCCGGATTTACCG CAAGGACCGTAATCATCCGATCTGCGTTACCGAATGGATGGATGAAT GCCGCCGCGAACCATTCAAAACTCGCGAAGGCAGAGAAATCACGGG GCCGTGGCAGTCGCATCCCAAACGGATGTTACGTCATAAAGCCATGA TTCAGTGTGCCCGTCTGGCCTTCGGATTTGCTGGTATCTATGACAAGG ATGAAGCCGAGCGCATTGTCGAAAATACTGCATACACTGCAGAACG TCAGCCGGAACGCGACATCACTCCGGTTAACGATGAAACCATGCAG GAGATTAACACTCTGCTGATCGCCCTGGATAAAACATGGGATGACGA CTTATTGCCGCTCTGTTCCCAGATATTTCGCCGCGACATTCGTGCATC GTCAGAACTGACACAGGCCGAAGCAGTAAAAGCTCTTGGATTCCTG AAACAGAAAGCCGCAGAGCAGAAGGTGGCAGCATGA |
| PapRecT | ATGGGTACTGCTCTAACGCCGTTATTAACCAAGTTTGCCACCCGCTA TGAGATGGGAACTACCCCCGAAGAGGTCGCTAATACGCTGAAACAG ACTTGTTTCAAGGGCCAAGTGAACGATAGTCAGATGGTAGCCCTTTT GATCGTTGCGGATCAATATAAGCTCAATCCATTCACAAAAGAGCTCT |

| | |
|---|---|
| | ACGCGTTCCCTGACAAAAATAATGGTATTGTTCCAGTTGTGGGAGTC<br>GATGGTTGGGCTAGAATTATTAACGAGAATCCCCAGTTTGATGGGAT<br>GGAATTCAGTATGGATCAACAGGGAACTGAATGCACTTGTAAAATTT<br>ACCGCAAAGACCGCTCGCACGCCATCAGCGCCACCGAGTACATGGC<br>TGAGTGCAAAAGGAACACTCAACCTTGGCAGTCTCACCCGCGACGTA<br>TGCTGCGTCATAAGGCTATGATTCAATGCGCCAGACTAGCCTTTGGT<br>TTCGCGGGGATCTACGATCAGGATGAGGCCGAACGCATTGTTGAACG<br>AGATGTAACTCCCGCCGAGCAATACGAGGATGTATCCGAAGCGATTT<br>GTCTGATCAAAGACTCACCAACTATGGAGGACTTGCAGGCCGCGTTC<br>TCAAACGCGTGGAAAGCTTACAAGACTAAAGGTGCCCGTGATCAAC<br>TGACTGCTGCTAAAGACCAGAGAAAAAAGGAGCTGTTGGATGCGCC<br>CATTGATGTCGAATTCGAAGAAACTGGAGATGATCGTGCTGCGTAA |
| MspRecT | ATGGCCGAGAATGCCGTCACGAAACAGGATTCCCCTAAGGCACCGG<br>AAACCATTAGTCAAGTGCTTCAGGTGCTGGTCCCACAATTGGCTCGT<br>GCAGTACCTAAAGGCATGGATCCTGATCGTATTGCACGTATCGTACA<br>GACGGAGATTCGCAAATCCCGCAACGCAAAAGCCGCTGGAATCGCA<br>AAGCAAAGTTTAGACGATTGCACACAGGAGTCCTTTGCGGGAGCCTT<br>ACTGACCTCAGCGGCTTTAGGGTTAGAGCCAGGCGTCAATGGGGAGT<br>GTTATCTGGTACCCTATCGTGATACACGCCGTGGTGTGGTCGAGTGC<br>CAACTGATTATTGGATATCAAGGGATTGTCAAACTTTTTTGGCAACA<br>TCCGCGCGCGAGCCGCATCGATGCGCAATGGGTTGGCGCGAACGAC<br>GAGTTCCATTATACGATGGGACTTAATCCTACCTTGAAACATGTAAA<br>GGCAAAAGGTGATCGTGGAAACCCGGTCTACTTTTACGCCATCGTCG<br>AGGTGACCGGTGCTGAGCCCTTATGGGATGTTTTTACTGCTGATGAA<br>ATTCGTGAACTTCGTCGTGGCAAGGTTGGATCGTCTGGAGATATTAA<br>GGACCCCCAGCGTTGGATGGAACGCAAGACAGCATTGAAACAGGTA<br>CTGAAATTGGCTCCCAAAACCACACGCCTGGATGCGGCGATCCGCGC<br>TGATGATCGTCCAGGGACTGACCTTTCACAGTCGCAGGCTCTGGCCT<br>TACCGTCTACCGTTAAGCCTACCGCAGACTATATTGATGGGGAGATC<br>GCCGAACCGCATGAAGTCGATACACCACCAAAGAGTTCACGTGCTC<br>AACGCGCACAACGTGCCACGGCACCGGCTCCTGATGTGCAAATGGC<br>CAACCCCGACCAATTGAAGCGTCTGGGGGAGATCCAAAAGGCGGAG<br>AAGTACAATGATGCCGACTGGTTCAAGTTTTTGGCGGACTCCGCCGG<br>TGTGAAAGCGACGCGTGCTGCTGATCTTACGTTTGATGAAGCAAAGG<br>CTGTAATCGACATGTTTGATGGGCCAAACGCGTGA |
| LrpRecT | ATGGCGAATCAAGTTGCACAGCAACAAAAACCGACAAAATTAACCG<br>ATCTGGTTTTGGATAGAGTCAAGCAGATGCAAGACACCCAGGACCTT<br>AGCCTTCCGAAAAACTATAACGCATCCAATGCACTGAATGCCGCGTT<br>TTTAGAATTGCAGAAGGTACAAGACCGGAACCACAGACCAGCACTG<br>GAAGTCTGCTCGCACGATTCTATTGTAAAATCGCTGTTGGACATGAC<br>TTTGCAGGGCTTATCCCCTGCGAAGGATCAGTGTTACTTCATAGTAT<br>ATGGCAATGAGTTACAGATGCAGAGATCTTATTCGGGACTGTCGCG<br>GCAGTTAAAAGATTAGATGGGGTGAAGAAGGTCCGGGCGGAAGTCG<br>TGCATGAAAAGGACGACTTCGAAATTGGCGCCAATGAAGACATGGA<br>GCTGGTAGTGAAACGGTTTGTACCAAAGTTCGAAAATCAAGACAAC<br>CAAATAATAGGGGCGTTCGCAATGATTAAAACGGATGAAGGTACGG |

| | |
|---|---|
| | ACTTCACAGTTATGACGAAAAAGGAAATAGATCAAAGTTGGGCGCA<br>AACACGCCAGAAGAACAATAAGGTACAGCAGAACTTTAGTCAAGAA<br>ATGGCGAAACGTACAGTCCTTAATCGTGCCGCTAAGATGTTTATAAA<br>CACTTCAGACGATTCGGACTTATTAACCGGGGCCATAAATGACACGA<br>CCTCAAACGAGTATGACGATGAAAGAAGAGATGTGACACCAGTCGA<br>GGACGAAAAACAGAGCACGGATAAATTACTGGAGGGGTTTCAGAAG<br>TCGCAGGAGGCGAAAGCAAAAGGGGTTAGTAACGACGGAAACAGTA<br>ATGAGGGAAAAGAGACAAGCGAGGAGGTGGCCGATGGACAGACGG<br>AACTGTTCTCTGAAGGTACTATTAAACCAGCAGATGAAGCGGATAGC<br>TAA |

- *M. smegmatis*

| | |
|---|---|
| MspRecT | ATGGCAGAAAACGCTGTAACCAAGCAAGACAGTCCCAAAGCGCCCG<br>AGACCATATCGCAGGTATTGCAAGTGTTAGTGCCTCAATTAGCAAGA<br>GCAGTCCCCAAAGGGATGGATCCTGACAGAATAGCACGCATAGTGC<br>AGACCGAAATACGTAAGTCCCGTAATGCCAAAGCTGCCGGCATCGC<br>AAAACAATCGTTAGATGATTGTACCCAGGAGAGTTTTGCCGGGGCGC<br>TGCTTACCTCAGCAGCCTTAGGTCTGGAACCAGGAGTTAACGGAGAG<br>TGTTATTTGGTCCCATACCGGGATACTCGTCGCGGAGTTGTTGAGTG<br>CCAACTTATTATCGGTTACCAGGGAATAGTGAAGTTGTTCTGGCAAC<br>ACCCTCGTGCGTCCCGGATTGACGCGCAGTGGGTAGGTGCAAACGAC<br>GAATTCCACTACACTATGGGCCTTAATCCGACACTTAAACACGTCAA<br>AGCGAAAGGGGATAGAGGAAACCCGGTGTACTTTTATGCAATTGTTG<br>AGGTTACTGGAGCAGAGCCGTTATGGGATGTCTTTACTGCCGATGAG<br>ATACGCGAGCTGCGTCGTGGCAAGGTCGGGAGTTCAGGGGACATTA<br>AAGATCCCCAACGGTGGATGGAGCGGAAGACTGCGCTGAAACAGGT<br>GTTGAAGTTGGCCCCCAAAACGACCCGCCTTGACGCTGCAATCCGGG<br>CGGATGATCGTCCTGGGACTGACCTGTCCCAAAGCCAAGCCTTAGCC<br>CTTCCAAGTACTGTCAAGCCAACCGCAGATTACATTGACGGGGAAAT<br>CGCAGAACCGCACGAAGTTGACACTCCTCCGAAGAGTAGCCGCGCA<br>CAACGTGCCCAGCGCGCGACGGCACCAGCGCCGGATGTGCAGATGG<br>CAAATCCTGACCAACTTAAAAGACTGGGAGAGATACAGAAAGCAGA<br>GAAGTACAACGACGCAGATTGGTTTAAGTTTTTGGCGGACAGCGCTG<br>GCGTCAAAGCAACTCGTGCGGCCGACTTGACCTTTGACGAAGCGAA<br>GGCGGTCATAGATATGTTTGATGGTCCAAACGCCTGA |
| PapRecT | ATGGGCACCGCCCTGACCCCACTCTTGACCAAGTTTGCCACGCGGTA<br>TGAGATGGGCACCACCCCAGAGGAAGTGGCGAACACCCTCAAGCAG<br>ACCTGCTTTAAGGGTCAGGTCAATGATAGCCAGATGGTGGCCTTGCT<br>GATCGTCGCGGACCAATATAAACTGAATCCATTTACCAAGGAACTCT<br>ATGCGTTTCCGGATAAGAACAATGGTATTGTCCCCGTCGTCGGCGTG<br>GACGGTTGGGCGCGGATCATTAACGAGAACCCCCAATTCGATGGCAT<br>GGAATTTTCGATGGACCAGCAAGGGACCGAATGCACCTGCAAAATC<br>TACCGGAAAGACCGTAGCCATGCCATTAGCGCCACGGAGTATATGG<br>CCGAATGTAAACGTAATACGCAGCCATGGCAATCCCATCCACGCCGG<br>ATGTTGCGCCACAAGGCGATGATCCAGTGTGCGCGGTTGGCCTTTGG<br>TTTCGCGGGCATCTATGACCAGGACGAAGCGGAACGCATCGTCGAG |

| | |
|---|---|
| | CGGGATGTGACCCCGGCCGAACAGTATGAGGACGTGTCGGAGGCGA TTTGTCTCATCAAAGATAGCCCAACGATGGAGGATTTGCAGGCCGCC TTCAGCAACGCCTGGAAGGCGTACAAGACCAAAGGTGCCCGTGACC AACTGACGGCCGCGAAGGACCAGCGTAAGAAGAACTGTTGGATGC CCCAATTGATGTCGAATTTGAGGAAACCGGGGACGATCGGGCGGCG TAA |

- *C. crescentus*

| | |
|---|---|
| λ Beta | ATGAGCACGGCGCTCGCGACGCTCGCGGGGAAGCTGGCCGAGCGTG TGGGCATGGATTCGGTCGATCCGCAGGAGCTCATCACCACGCTCCGG CAGACGGCCTTTAAGTGTGACGCGAGCGATGCCCAGTTTATCGCCCT CCTGATCGTGGCCAATCAGTACGGCCTGAACCCGTGGACGAAGGAA ATCTACGCCTTTCCCGACAAGCAAAACGGGATCGTGCCGGTGGTCGG CGTCGATGGGTGGTCCCGTATCATCAATGAAAATCAGCAATTTGATG GCATGGATTTCGAGCAAGACAATGAATCCTGCACGTGCCGCATCTAT CGGAAGGACCGCAACCATCCGATCTGCGTGACGGAATGGATGGATG AGTGCCGCCGGGAGCCCTTTAAGACGCGGGAGGGCCGGGAAATCAC CGGGCCCTGGCAGTCGCACCCCAAGCGGATGCTCCGTCATAAGGCG ATGATCCAATGTGCCCGCCTCGCCTTCGGGTTCGCGGGCATCTACGA CAAGGATGAAGCCGAGCGCATCGTGGAAAATACGGCCTACACGGCG GAGCGTCAGCCGGAACGGGATATCACGCCGGTCAATGACGAAACGA TGCAGGAAATCAATACCCTGCTCATCGCGCTCGACAAGACCTGGGAC GATGATCTGCTGCCCCTGTGTAGCCAAATCTTCCGTCGTGATATCCG CGCCTCGTCCGAACTGACCCAAGCGGAGGCGGTGAAGGCCCTGGGG TTCCTGAAGCAGAAGGCCACCGAGCAAAAGGTCGCGGCCTAA |
| PapRecT | ATGGGCACGGCGCTCACGCCGCTGCTCACCAAGTTTGCCACCCGTTA CGAGATGGGGACCACCCCCGAAGAAGTGGCGAACACCCTGAAGCAA ACGTGCTTCAAGGGCCAGGTCAACGACTCGCAGATGGTGGCCCTGCT CATCGTGGCCGATCAGTATAAGCTCAATCCGTTCACCAAGGAACTCT ACGCGTTTCCCGATAAGAACAATGGGATCGTGCCGGTCGTCGGCGTC GACGGCTGGGCGCGTATCATCAATGAAAATCCGCAGTTCGACGGCA TGGAATTCTCGATGGACCAACAAGGGACCGAATGTACGTGCAAGAT CTATCGTAAGGATCGTTCGCACGCGATCAGCGCCACGGAATACATGG CGGAGTGTAAGCGGAATACGCAGCCGTGGCAATCCCACCCCCGCCG TATGCTGCGCCATAAGGCGATGATCCAATGTGCCCGCCTGGCGTTTG GGTTCGCCGGCATCTACGATCAAGATGAAGCGGAGCGGATCGTCGA ACGCGATGTGACGCCCGCCGAACAATATGAAGACGTGTCGGAAGCG ATCTGCCTGATCAAGGACAGCCCCACGATGGAAGATCTCCAAGCGG CCTTTAGCAATGCCTGGAAGGCCTACAAGACGAAGGGGGCGCGTGA CCAACTGACGGCGGCCAAGGATCAACGGAAGAAGGAGCTGCTGGAT GCGCCGATCGATGTCGAATTCGAGGAAACGGGGGACGATCGTGCCG CGTAA |

Nucleotide sequences of codon optimized Genes: SSB

- *L. lactis*

| | |
|---|---|
| EcSSB | ATGGCAAGCCGTGGGGTTAACAAAGTTATTCTTGTTGGAAACTTAGG<br>ACAAGACCCGGAAGTGCGTTATATGCCTAATGGAGGCGCGGTAGCC<br>AATATCACCTTGGCCACAAGCGAGTCTTGGCGAGACAAAGCAACAG<br>GTGAAATGAAAGAACAAACTGAATGGCACAGAGTAGTTTTGTTTGG<br>AAAATTGGCAGAGGTAGCCTCAGAATACTTGCGAAAGGGCAGTCAG<br>GTCTATATAGAGGGCCAATTGCGTACCCGTAAGTGGACAGACCAGA<br>GCGGACAAGATCGTTATACGACCGAGGTCGTTGTTAATGTAGGAGG<br>CACAATGCAGATGTTGGGGGGGAGACAGGGCGGAGGCGCTCCGGCT<br>GGAGGCAATATCGGGGGTGGCCAACCTCAAGGTGGGTGGGGGCAGC<br>CACAGCAACCGCAAGGAGGTAATCAATTTAGTGGAGGAGCCCAATC<br>ACGTCCGCAGCAGTCTGCGCCTGCCGCCCCTTCTAATGAACCGCCGA<br>TGGATTTTGACGATGATATACCTTTCTGA |
| PaSSB | ATGGCCCGTGGAGTGAACAAAGTAATTCTTGTCGGTAATGTGGGTG<br>GGGATCCAGAGACGCGATACATGCCAAACGGGAACGCCGTGACAAA<br>TATCACCTTAGCCACGAGCGAATCTTGGAAGGACAAACAAACAGGT<br>CAGCAACAAGAACGAACCGAATGGCATAGAGTTGTATTTTTTGGCC<br>GACTTGCTGAGATCGCGGGTGAGTACCTTAGAAAGGGTTCTCAGGTT<br>TATGTCGAGGGCTCATTAAGAACACGTAAGTGGCAGGGGCAGGACG<br>GGCAAGACCGATATACAACTGAAATAGTAGTGGACATAAACGGCAA<br>CATGCAACTTCTTGGTGGCAGACCGAGTGGGGACGATTCACAGAGA<br>GCTCCAAGAGAACCTATGCAGCGACCACAGCAGGCTCCTCAACAGC<br>AGTCTCGTCCGGCCCCTCAGCAGCAACCGGCTCCGCAACCTGCACAA<br>GATTACGATAGTTTTGATGATGATATTCCATTCTAA |
| MsSSB | ATGGCGGGAGACACAACAATTACGGTTGTGGGAAACTTGACAGCCG<br>ATCCTGAATTGCGATTCACCCCATCAGGCGCTGCGGTGGCGAATTTC<br>ACAGTCGCGAGCACCCCACGAATGTTTGATAGACAATCAGGCGAAT<br>GGAAGGATGGCGAAGCGTTGTTTTTACGATGCAACATCTGGAGAGA<br>GGCGGCCGAGAACGTCGCCGAAAGCCTTACCCGTGGCAGTCGAGTG<br>ATTGTAACCGGACGATTAAAGCAAAGAAGTTTTGAGACGAGAGAAG<br>GAGAGAAACGAACTGTGGTAGAGGTAGAGGTGGATGAAATAGGTCC<br>TAGTTTGCGTTATGCCACAGCGAAAGTAAACAAAGCCTCTCGTAGTG<br>GTGGCGGGGGGGGCGGCTTTGGTTCAGGGGGTGGAGGTTCACGACA<br>GAGCGAGCCAAAGGATGATCCTTGGGGCAGTGCCCCTGCATCAGGC<br>AGTTTTAGCGGAGCAGATGACGAGCCGCCTTTTTGA |
| LrSSB | ATGCTTAATCGTGCAGTCTTAACTGGGCGTTTAACAAGAGATCCCGA<br>GTTGCGGTACACAACCAGCGGGACAGCAGTTGTTTCATTTACGTTAG<br>CTGTTGATCGGCAATTCCGAAACCAAAATGGTGATCGTGATGCTGAT<br>TTTATCAATTGCGTTATTTGGCGTAAATCCGCTGAAAACTTTAGTAA<br>CTTTACGCATAAGGGTTCACTTGTTGGAATTGAAGGGCGTATTCAAA<br>CCCGGAATTATGAAAACCAACAGGGTAACCGTGTGTATGTTACCGA<br>AGTTGTTGTAGATAATTTTGCATTGTTAGAACCTCGTCAAAATGGTG<br>GCATGAACCAATCAGGGGTTCAACAACCATTTAACAGCAACCAACA<br>ATCATTTGGTGCTCAGGCTCCACAATATGGTAGTCAACCACAACCTG<br>GAAATAATGCTCCTCAAAGTAATCCGTCACCAAGTATGGATAATGGT<br>TTCGATCCCAATCAAAATGCTGGCAACCAGTTCCCTGGAAGCAGTGA |

| | |
|---|---|
| | TGATGGTGGTCAATCCATTGATTTAGCTGATGACGAATTACCATTCT AA |
| LlSSB | ATGATTAACAATGTTGTATTAGTGGGACGCATTACTCGCGATCCTGA ACTTCGTTACACCCCTCAAAATCAAGCTGTTGCAACTTTTTCATTGGC TGTAAATCGTCAATTTAAAAATGCTAACGGTGAACGTGAGGCTGATT TCATTAACTGCGTTATTTGGCGCCAACAAGCTGAAAATTTGGCAAAT TGGGCTAAAAAAGGAGCTTTGATCGGTGTAACTGGTCGAATTCAAA CACGTAATTATGAAAATCAACAAGGTCAACGCGTTTATGTGACTGA GGTTGTGGCTGATAGTTTCCAAATGTTGGAAAGTAGATCTGCTCGCG ATGGTATGGGAGGCGGAGCTTCTGCCGGTTCATATTCTGCACCAAGC CAATCTACAAATAATACTCCACGTCCACAAACGAATAATAATAGTG CAACACCGAATTTCGGTCGTGATGCTGACCCATTTGGTAGCTCACCT ATGGAAATCTCGGATGATGATCTTCCATTCTAA |
| PapSSB | ATGCGTGGGGTTAATAAGGTAATCTTAGTTGGTAACGTGGGTGGGG ACCCGGAGACCCGATATATGCCAAATGGAAACGCGGTAACAAACAT CACCCTTGCAACTAGTGAGAGTTGGAAAGATAAACAAACTGGCCAA CAGCAAGAACGTACTGAATGGCACAGAGTGGTGTTTTTGGCAAATT AGCCGAAATTGTCGGCCAACACGTTAAGAAAGGCCAGCAGCTTTAC GTCGAAGGGTCATTGCGAACCCGTAAGTGGCAAGCGCAGGATGGTC AGGACAGATATACGACAGAAATCATAGTAGATATGCACGGACAAAT GCAAATGTTCGGGGGAAAACCTGGGAATGAGCAGGCCGCACAGTCA AGATCATCTACCCAACAACAAAGCGCCCCGCAACAACGATCAGCAC AGGATGAATTTGATGATGATATACCTTTATAA |

- *E. coli*

| | |
|---|---|
| EcSSB | ATGGCCAGCAGAGGCGTAAACAAGGTTATTCTCGTTGGTAATCTGG GTCAGGACCCGGAAGTACGCTACATGCCAAATGGTGGCGCAGTTGC CAACATTACGCTGGCTACTTCCGAATCCTGGCGTGATAAAGCGACCG GCGAGATGAAAGAACAGACTGAATGGCACCGCGTTGTGCTGTTCGG CAAACTGGCAGAAGTGGCGAGCGAATATCTGCGTAAAGGTTCTCAG GTTTATATCGAAGGTCAGCTGCGTACCCGTAAATGGACCGATCAATC CGGTCAGGATCGCTACACCACAGAAGTCGTGGTGAACGTTGGCGGC ACCATGCAGATGCTGGGTGGTCGTCAGGGTGGTGGCGCTCCGGCAG GTGGCAATATCGGTGGTGGTCAGCCGCAGGGCGGTTGGGGTCAGCC TCAGCAGCCGCAGGGTGGCAATCAGTTCAGCGGCGGCGCGCAGTCT CGCCCGCAGCAGTCCGCTCCGGCAGCGCCGTCTAACGAGCCGCCGA TGGACTTTGATGATGACATTCCGTTCTGA |
| PaSSB | ATGGCTCGCGGGGTAAATAAGGTCATTTTGGTTGGCAATGTTGGTGG TGATCCCGAGACACGCTATATGCCTAACGGGAACGCCGTCACTAAT ATCACACTGGCAACGTCCGAGTCATGGAAGGATAAACAGACAGGTC AACAGCAAGAACGCACGGAGTGGCACCGCGTGGTATTTTTCGGGCG TCTTGCTGAGATCGCCGGAGAGTATTTACGCAAAGGATCGCAGGTAT ACGTTGAGGGTTCTTTACGCACGCGCAAGTGGCAGGGTCAGGATGG TCAGGACCGTTATACTACCGAAATTGTAGTCGACATTAACGGGAAC ATGCAATTATTAGGTGGTCGTCCGAGCGGAGATGACTCCCAGCGCG CCCCCCGCGAGCCCATGCAGCGTCCGCAACAGGCTCCACAGCAGCA |

| | |
|---|---|
| | GAGCCGCCCTGCCCCTCAACAACAACCCGCTCCTCAACCCGCGCAA<br>GATTACGATTCGTTTGACGACGATATTCCTTTTTAA |
| MsSSB | ATGGCAGGGGATACCACGATAACCGTTGTCGGTAACTTAACCGCGG<br>ACCCTGAACTTCGTTTCACACCATCCGGTGCAGCGGTTGCAAACTTC<br>ACGGTCGCTTCTACGCCTCGTATGTTCGACAGACAGTCTGGTGAGTG<br>GAAAGATGGGGAAGCACTGTTTTTAAGATGCAATATATGGCGCGAA<br>GCAGCAGAGAATGTAGCCGAGAGTTTAACCAGAGGTTCACGTGTGA<br>TCGTAACTGGCCGTTTGAAACAACGCTCCTTTGAAACACGCGAAGGC<br>GAGAAACGCACGGTAGTTGAGGTCGAAGTCGACGAGATAGGCCCGT<br>CCTTACGCTATGCCACAGCGAAAGTCAACAAAGCGTCTCGCAGCGG<br>AGGCGGTGGGGGCGGGTTTGGTAGTGGTGGGGGGGGTAGTCGTCAA<br>TCGGAACCCAAGGATGACCCGTGGGGGTCGGCACCAGCTTCAGGAA<br>GTTTTTCTGGGGCCGATGACGAGCCGCCATTTTGA |
| LrSSB | ATGCTGAACCGTGCCGTGCTTACTGGTCGCCTTACTCGTGACCCTGA<br>ATTGCGCTATACGACATCAGGGACTGCAGTAGTGTCCTTTACATTGG<br>CGGTCGATCGTCAATTTCGTAACCAAAACGGCGACCGCGACGCCGA<br>TTTTATCAACTGTGTGATTTGGAGAAAGAGCGCCGAGAACTTTAGCA<br>ATTTCACTCATAAAGGGAGTTTAGTTGGAATCGAGGGGCGTATCCAA<br>ACGAGAAACTACGAAAACCAGCAAGGCAATCGCGTCTACGTAACCG<br>AAGTCGTAGTAGATAACTTCGCCCTGTTGGAACCACGGCAAAACGG<br>TGGGATGAACCAATCTGGAGTTCAACAACCCTTCAACAGTAACCAG<br>CAGTCTTTCGGGGCTCAGGCACCTCAATATGGCAGTCAGCCACAACC<br>TGGAAACAATGCCCCACAGTCTAACCCAAGTCCCTCTATGGACAATG<br>GGTTTGACCCCAACCAGAATGCGGGGAACCAATTCCCTGGGAGCTC<br>GGATGACGGCGGCCAATCAATTGATCTGGCTGACGATGAATTACCCT<br>TTTAA |

- *M. smegmatis*

| | |
|---|---|
| PaSSB | ATGGCGCGTGGGGTGAACAAAGTCATCCTCGTGGGGAATGTCGGTG<br>GCGATCCCGAAACGCGTTATATGCCGAATGGGAATGCGGTCACCAA<br>TATCACGCTCGCCACCAGCGAGTCCTGGAAAGATAAACAAACGGGT<br>CAACAGCAGGAGCGTACGGAGTGGCATCGGGTGGTCTTCTTCGGGC<br>GCCTCGCCGAGATCGCCGGGGAATACCTCCGTAAAGGTTCGCAGGT<br>CTATGTGGAGGGCTCGCTGCGGACCCGTAAATGGCAAGGTCAGGAT<br>GGCCAGGATCGGTACACGACGGAAATCGTCGTGGACATTAACGGTA<br>ATATGCAATTGCTCGGTGGCCGCCCCTCCGGCGATGATAGCCAGCGT<br>GCCCCGCGTGAACCGATGCAACGCCCGCAACAAGCGCCCCAACAGC<br>AATCGCGGCCCGCGCCGCAGCAGCAGCCGGCCCCGCAACCAGCCCA<br>GGACTACGATTCGTTTGATGATGACATTCCATTTTAA |

- *C. crescentus*

| | |
|---|---|
| PaSSB | ATGGCGCGTGGGGTCAATAAGGTGATCCTGGTCGGCAACGTGGGGG<br>GCGATCCCGAAACCCGGTACATGCCGAACGGCAACGCGGTCACCAA<br>CATCACCCTGGCGACCAGCGAGAGCTGGAAGGATAAGCAAACGGGC<br>CAGCAGCAAGAACGTACGGAATGGCATCGTGTGGTCTTTTTCGGCC<br>GGCTGGCGGAGATCGCGGGGGAATACCTCCGTAAGGGGTCCCAAGT |

| | |
|---|---|
| | CTACGTGGAGGGCTCGCTGCGGACCCGGAAGTGGCAAGGGCAAGAT GGGCAAGATCGCTACACCACGGAGATCGTCGTCGACATCAACGGGA ACATGCAGCTCCTCGGGGGGCGTCCCTCGGGGGACGATTCCCAACG CGCCCCCCGTGAGCCCATGCAACGCCCGCAGCAAGCGCCCCAGCAA CAATCGCGTCCCGCCCCCCAGCAACAGCCGGCGCCCCAACCGGCGC AGGACTACGACTCGTTCGACGATGATATCCCCTTTTAA |

Nucleotide sequences of codon optimized Genes: phage exonuclease

- *L. lactis*

| | |
|---|---|
| PapExo | ATGATAGAACAGCGTAGTGATGAATGGTTCGCGCAGCGACTTGGCC GAGTCACCGCGAGTAAAGTAAAGGATGTCATGGCGAAGGGGCGATC AGGTGCGCCATCAGCCACCAGACAGAATTACATGATGCAATTGTTAT GTGAGAGACTTACCGGGAAACGAGAAGAGGGGTTCACGAGTGCGGC GATGCAGCGTGGGACGGACCTTGAACCAATAGCGCGATCAGCTTAT GAGTTTAACGCAGGAGTAATGACTATAGAAACAGGCCTTATTATCC ATCCACGTATCGACGGTTTCGGAGCTAGTCCGGATGGGCTTGCGGGA GAGCATGGATTAGTGGAAATTAAGTGCCCGTCAACAGCAACGCACA TTTATACCATGCAAAGTGGTAAGCACGACCCTCAGTACGAATGGCA AATGCTTGCTCAAATGAGTTGCTCAGGCAGAGAGTGGGTGGATTTCG TGTCATTCGACGATAGATTGCCAGACGAATTGCAATATGTTTGTTTC CGTTATCACCGTGATGAAGAGAGAATAAGAGAAATGGAAAGCGAA GTTAAGGCATTCTTGGAGGAATTAGCTGAATTGGAACACCAAATGC GTGAACGTATGAGAAAGGCGGCCTAA |
| LrpExo | ATGAAACTTACGGCCAACAATTACTATAGCCATGAGACTGACTGGC AATATATGTCAGTTTCATTGTTCAAAGACTTCGAAAAGTGCGAAGCG CGTGCATTAGCAAAGTTGAAGGAAGATTGGCAACCTGTTTCTAGTCC AGTTCCGCTTTTGGTTGGGAACTATGTACACAGTTATTTCGAAAGTG CTAAGAGCCACCAAGATTTTATAGAGGCGAATAAGAAAGAGCTTAT GACCAGACCTACTAAGACAAACCCGAACGGCCATCTTAGAGCGGAA TTTAAGGGGGCAAACTCAATGATTCAGACCTTGCAAGCCGACGATA TGTTTAACTACTTTTATGCACCAGGGGACAAAGAAGTTATCGTTACC GGAGAGATAGACGGCTATTTGTGGAAGGGAAAAATAGACTCTTTAG TTCTTGACAAAGGCTATTTTTGCGATCTTAAGACGGTAGACGACATT CATAAGGGACATTGGAATACGTATGAACACAGATACGTCCCGTTCA TTCAAGACCGAGAATATGATTTACAAATGGCTGTTTATAGAGAGTTA ATCAAGCAGACGTTCGGGAAAAAGTGCCAACCTTTAATTTTTGCCAT CTCTAAGCAAACTCCGCCTGACAAGATGGCCATCGACTTTAATGGCG TTGATGACGACTATCAGATGCAGGCCGATCTTGATAAGGTCAAAGA GCTTCAACCACACTTTTGGAAAGTAATGACGGGAGAGGAAGAGCCT GTCCACTGTGGTAAGTGCGACTATTGTAGAGAAACGAAAATGTTGA GCGGCTTCATCCACGCATCAGAAATAGAGGTTTAA |

Negative control (no recombinase)

- All organisms

| | |
|---|---|
| mCardinal **RBS** eGFP | ATGCACCATCATCACCACCACGGTTCCGGCATGGTTTCTAAAGGTGA AGAACTGATCAAGGAAAACATGCACATGAAGCTGTATATGGAAGGT |

| | |
|---|---|
| | ACCGTTAACAACCACCATTTCAAATGCACCACTGAAGGTGAAGGTAA ACCGTACGAGGGTACGCAGACCCAACGTATTAAAGTTGTTGAGGGT GGTCCGCTGCCGTTCGCGTTCGACATCCTGGCGACCTGTTTCATGTAC GGCTCTAAAACCTTCATCAACCACACCCAGGGTATCCCTGACTTCTT CAAACAGTCTTTCCCGGAGGGTTTCACCTGGGAACGTGTTACCACCT ACGAAGACGGTGGTGTACTGACCGTTACCCAGGACACTTCTCTGCAG GACGGTTGCCTGATCTACAACGTTAAACTCCGCGGTGTTAATTTCCC GTCTAACGGTCCGGTTATGCAAAAAAAGACGCTGGGTTGGGAAGCG ACTACGGAAACTCTCTACCCTGCCGATGGCGGCCTCGAAGGTCGTTG TGATATGGCGCTGAAACTGGTTGGTGGCGGTCACCTGCACTGCAATC TGAAAACTACCTACCGTTCTAAAAAACCAGCTAAAAACCTCAAAATG CCGGGTGTTTACTTTGTTGATCGTCGTCTGGAACGTATCAAAGAAGC AGACAACGAAACTTACGTTGAACAGCACGAAGTTGCGGTGGCGCGT TACTGCGACCTGCCATCTAAACTGGGTCACAAAGGTATGGACGAACT GTACAAATAA**AAAAAATAGGAGGAAAAACAT**ATGGGTTCTCACCA CCATCACCACCACAGCGGCTCTAAAGGTGAAGAATTATTCACTGGTG TTGTCCCAATTTTGGTTGAATTAGATGGTGATGTTAATGGTCACAAAT TTTCTGTCTCCGGTGAAGGTGAAGGTGATGCTACGTACGGTAAATTG ACCTTAAAATTTATTTGTACTACTGGTAAATTGCCAGTTCCATGGCCA ACCTTAGTCACTACTTTCACTTATGGTGTTCAATGTTTTTCTAGATAC CCAGATCATATGAAACAACATGACTTTTTCAAGTCTGCCATGCCAGA AGGTTATGTTCAAGAAAGAACTATTTTTTTCAAAGATGACGGTAACT ACAAGACCAGAGCTGAAGTCAAGTTTGAAGGTGATACCTTAGTTAAT AGAATCGAATTAAAAGGTATTGATTTTAAAGAAGATGGTAACATTTT AGGTCACAAATTGGAATACAACTATAACTCTCACAATGTTTACATCA TGGCTGACAAACAAAAGAATGGTATCAAAGTTAACTTCAAAATTAG ACACAACATTGAAGATGGTTCTGTTCAATTAGCTGACCATTATCAAC AAAATACTCCAATTGGTGATGGTCCAGTCTTGTTACCAGACAACCAT TACTTATCCACTCAATCTGCCTTATCCAAAGATCCAAACGAAAAGAG AGACCACATGGTCTTGTTAGAATTTGTTACTGCTGCTGGTATTACCCA TGGTATGGATGAATTGTACAAATAA |

<u>Nucleotide sequences of codon optimized Genes: dominant negative mutL proteins</u>

- *L. lactis*

| | |
|---|---|
| E. coli MutL E32K | ATGCCTATACAAGTGTTGCCTCCACAGTTGGCCAACCAAATCGCG GCAGGCGAGGTGGTCGAACGTCCGGCTTCAGTCGTTAAGGAATTG GTAAAAAATTCTTTGGATGCAGGGGCAACGAGAATTGATATTGAC ATCGAACGAGGCGGGGCCAAGTTAATCAGAATCCGAGACAATGG GTGTGGGATTAAAAAGGATGAACTTGCTTTGGCGTTGGCACGTCA CGCGACCAGCAAAATAGCGTCTCTTGACGACTTGGAAGCTATTAT CAGTCTTGGTTTCCGTGGGGAAGCCTTAGCATCTATTAGCTCTGTG TCACGTTTGACTTTGACTAGCAGAACGGCGGAACAGCAGGAAGC ATGGCAAGCGTATGCGGAAGGACGAGACATGAACGTCACGGTTA AGCCGGCAGCCCACCCGGTCGGCACGACCTTGGAGGTCTTGGACT TGTTCTATAATACCCCTGCACGTCGTAAATTCTTACGAACCGAAA AGACCGAATTTAACCATATAGATGAGATAATAAGAAGAATTGCG |

| | |
|---|---|
| | TTAGCACGTTTCGATGTTACTATAAATTTGAGTCATAACGGAAAA<br>ATCGTTAGACAGTATCGAGCCGTGCCTGAGGGCGGGCAGAAGGA<br>AAGAAGATTAGGGGCTATTTGTGGCACTGCTTTTCTTGAACAAGC<br>ACTTGCGATCGAATGGCAACATGGGGACCTTACCTTGCGAGGTTG<br>GGTAGCGGACCCGAATCATACAACACCAGCGTTGGCAGAGATAC<br>AATATTGCTATGTAAACGGACGAATGATGAGAGATCGTTTGATCA<br>ACCACGCAATACGACAGGCTTGCGAAGATAAGTTGGGGGCGGAT<br>CAACAGCCAGCTTTCGTCCTTTATCTTGAAATTGACCCTCATCAGG<br>TAGATGTGAATGTACATCCGGCCAAACACGAGGTTCGTTTTCATC<br>AAAGTCGACTTGTGCATGATTTTATATACCAGGGTGTCTTAAGTG<br>TCTTGCAGCAGCAGCTTGAGACACCTTTACCTTTAGATGATGAGC<br>CGCAGCCAGCTCCGCGTAGTATCCCTGAGAATCGAGTTGCCGCCG<br>GCAGAAATCATTTCGCAGAACCGGCAGCCCGTGAACCTGTAGCAC<br>CGAGATACACCCCGGCTCCTGCCTCTGGATCACGTCCTGCTGCCC<br>CGTGGCCTAACGCACAACCGGGCTATCAGAAGCAGCAGGGTGAA<br>GTTTATCGTCAATTGTTACAAACTCCGGCACCAATGCAAAAACTT<br>AAGGCCCCGGAGCCGCAGGAACCGGCGCTTGCTGCAAATTCACA<br>ATCTTTCGGACGAGTTTTAACAATAGTGCATAGTGACTGCGCATT<br>ACTTGAGCGTGACGGCAACATTAGTTTGCTTTCATTGCCTGTTGCC<br>GAGCGTTGGTTGAGACAAGCACAATTAACCCCTGGTGAAGCACC<br>AGTCTGTGCACAGCCATTATTGATCCCATTGCGTTTAAAGGTCTCA<br>GCCGAGGAAAAGAGTGCTTTGGAAAAAGCCCAAAGTGCCCTTGC<br>AGAGCTTGGAATTGATTTCCAAAGCGACGCACAACACGTTACGAT<br>AAGAGCGGTTCCATTACCGTTAAGACAGCAAAACTTACAAATTCT<br>TATACCAGAGCTTATCGGGTATTTGGCGAAACAGAGCGTATTCGA<br>ACCAGGTAATATCGCCCAGTGGATAGCGCGTAACCTTATGTCAGA<br>ACACGCGCAGTGGAGTATGGCGCAAGCTATCACATTGTTAGCCGA<br>CGTTGAGCGTTTGTGCCCACAGTTGGTGAAAACGCCTCCGGGTGG<br>ACTTCTTCAAAGTGTGGACTTACATCCAGCAATTAAGGCTCTTAA<br>AGATGAATAA |
| L. lactis MutL E33K | GTGGGAAAAATTATTGAACTAAATGAAGCGCTCGCCAATCAAATT<br>GCTGCTGGAGAGGTGGTTGAGCGGCCTGCTAGTGTTGTCAAAGAA<br>TTAGTCAAAAACTCAATTGATGCTGGAAGCAGTAAAATTATTATC<br>AATGTTGAAGAAGCAGGTTTGCGATTAATTGAAGTCATTGATAAT<br>GGTTTGGGCTTAGAAAAAGAAGATGTGGCTTTGGCTTTGCGTCGT<br>CATGCGACAAGTAAAATCAAAGATTCAGCTGATTTATTTCGAATT<br>AGAACGCTCGGTTTTCGGGGTGAGGCTCTGCCGTCAATCGCTTCT<br>GTCAGTCAGATGACGATTGAAACAAGTAATGCTCAGGAAGAAGC<br>TGGGACAAAACTGATTGCTAAAGGTGGGACGATTGAAACTTTAG<br>AACCTCTTGCAAAGCGGTTAGGGACAAAAATTTCTGTTGCGAATC<br>TTTTTTATAATACACCAGCAAGGCTCAAGTATATCAAGTCTTTACA<br>GGCTGAACTTTCTCATATTACAGATATTATCAATCGTTTGAGCCTC<br>GCTCATCCAGAGATTTCTTTTACTTTAGTTAATGAGGGTAAAGAA<br>TTTTTGAAAACGGCGGGAAATGGAGACTTGCGCCAAGTGATTGCT<br>GCAATTTATGGCATTGGAACGGCGAAAAAAATGCGTGAGATTAA<br>TGGCTCGGACTTAGATTTTGAACTGACAGGTTATGTCAGTTTACCC |

| | |
|---|---|
| | GAGCTGACAAGAGCGAATCGCAACTATATCACGATTTTGATTAAT<br>GGTCGATTTATCAAGAATTTTTTGTTGAATCGAGCAATTTTAGAA<br>GGTTACGGGAACCGATTGATGGTTGGACGTTTTCCTTTTGCTGTTT<br>TATCAATTAAAATTGACCCTAAATTAGCAGATGTCAATGTCCATC<br>CGACAAAACAAGAAGTACGTTTGTCTAAGGAACGTGAATTGATG<br>ACTTTAATTTCTAAAGCGATTGATGAGACCTTATCAGAAGGGGTT<br>TTGATTCCAGAAGCTTTGGAAAATTTGCAAGGTAGAGCCAAGGAA<br>AAGGGGACTGTTTCTGTTCAAACGGAACTTCCTTTACAGAATAAT<br>CCTTTATACTATGACAATGTTCGTCAAGATTTTTTTGTCAGAGAAG<br>AAGCGATTTTTGAAATCAATAAAAACGATAATTCAGATTCTCTGA<br>CTGAACAAAATTCTACTGATTATACAGTTAATCAGCCAGAAACTG<br>GTTCTGTCAGTGAAAAAATTACGGACAGAACTGTCGAAAGTTCAA<br>ATGAATTTACTGACAGAACCCCAAAAAATTCTGTCAGTAACTTTG<br>GAGTTGATTTTGATAATATTGAGAAGCTGAGTCAGCAATCAACTT<br>TTCCCCAACTAGAATACTTGGCACAATTGCATGCGACTTATTTACT<br>TTGTCAGTCAAAAGAGGGTCTTTATTTGGTTGACCAACATGCGGC<br>TCAGGAGCGAATCAAGTATGAATATTGGAAAGATAAAATCGGCG<br>AAGTGAGCATGGAGCAACAAATTTTACTTGCGCCATATTTATTTA<br>CTTTACCCAAAAATGATTTTATTGTTTTAGCTGAGAAAAAGGATTT<br>ATTACATGAAGCAGGGGTTTTCTTGGAAGAATACGGAGAAAATC<br>AATTCATATTAAGAGAGCATCCGATTTGGTTAAAAGAAACTGAGA<br>TAGAGAAATCAATTAATGAAATGATTGATATTATTCTCTCATCAA<br>AAGAATTTTCACTCAAAAAATATCGGCATGATTTAGCCGCAATGG<br>TTGCTTGTAAAAGCTCAATCAAAGCCAACCATCCCCTTGATGCCG<br>AGTCTGCTAGAGCTTTGCTTAGAGAATTATCAACTTGTAAAAATC<br>CTTATAGTTGTGCGCATGGACGGCCAACGATTGTCCATTTTTCAG<br>GAGATGACATTCAAAAAATGTTCCGCAGAATTCAAGAAACGCAT<br>CGTTCAAAAGCGGCCTCTTGGAAAGATTTTGAGTAA |
| *L. lactis* dsDNA template (Erythromycin resistance gene)<br>Homology arm, promoter, gene, terminator, homology arm | ATGATTGAACTTAGTGGCAAAGATAGAAAGTATTTGTATAAACTA<br>GTAAAATCCAAAAAACTAAATTATGAACAAGGTAATTTATCGCAT<br>CAAGTTTTAATTGAAAACAAGTTAGCAAAAGTTTACTTTACAAGC<br>GATAAATATGATCCTGACTTAGGGGAACACATAAATCCACAAAAT<br>ATTATTGCTCCAACTAGTACAGGTTTAAGATATAAAAATATTTAT<br>CGTGAACAATTATGGGAAAAATATTTTACTCCTATTTGGGTATCT<br>ACGGCAACAACGACTCTAATATGGTTAGCAAAATATTTACTAGAG<br>AACTTGCTGTAACGCTAAGTAAGATTACTATCCATAGCTCTTTTTT<br>ATCTTTTCTCATCTTTCCACCTCCTAGCCCACTCGGGCTTTTTAATT<br>TAAAAATTGTTTAATCTCATGAAACGCCATGCCTATTTCTAACAGT<br>AAGATAATGCTGTCAGTATAGCGCCTAAGCGTTTCTTTTTGTTCTG<br>ATTTTTTAATGTGGTCTTTATTCTTCAACTAAAGCACCCATTAGTT<br>CAACAAACGAAAATTGGATAAAGTGGGATATTTTTAAAATATATA<br>TTTATGTTACAGTAATATTGACTTTTAAAAAAGGATTGATTCTAAT<br>GAAGAAAGCAGACAAGTAAGCCTCCTAAATTCACTTTAGATAAA<br>AATTTAGGAGGCATATCAAATGAACAAAAATATAAAATATTCTCA<br>AAACTTTTTAACGAGTGAAAAAGTACTCAACCAAATAATAAAAC<br>AATTGAATTTAAAAGAAACCGATACCGTTTACGAAATTGGAACAG |

| | |
|---|---|
| | GTAAAGGGCATTTAACGACGAAACTGGCTAAAATAAGTAAACAG<br>GTAACGTCTATTGAATTAGACAGTCATCTATTCAACTTATCGTCAG<br>AAAAATTAAAACTGAATACTCGTGTCACTTTAATTCACCAAGATA<br>TTCTACAGTTTCAATTCCCTAACAAACAGAGGTATAAAATTGTTG<br>GGAGTATTCCTTACCATTTAAGCACACAAATTATTAAAAAAGTGG<br>TTTTTGAAAGCCATGCGTCTGACATCTATCTGATTGTTGAAGAAG<br>GATTCTACAAGCGTACCTTGGATATTCACCGAACACTAGGGTTGC<br>TCTTGCACACTCAAGTCTCGATTCAGCAATTGCTTAAGCTGCCAG<br>CGGAATGCTTTCATCCTAAACCAAAAGTAAACAGTGTCTTAATAA<br>AACTTACCCGCCATACCACAGATGTTCCAGATAAATATTGGAAGC<br>TATATACGTACTTTGTTTCAAAATGGGTCAATCGAGAATATCGTC<br>AACTGTTTACTAAAAATCAGTTTCATCAAGCAATGAAACACGCCA<br>AAGTAAACAATTTAAGTACCGTTACTTATGAGCAAGTATTGTCTA<br>TTTTTAATAGTTATCTATTATTTAACGGGAGGAAATAATAATATG<br>AGATAATGCCGACTGTACTTTTTACAGTCGGTTTTCTAATGTCACT<br>AACCTGCCCCGTTAGTTGAAGAAGGTTTTTATATTACAGCTCCAC<br>GGTTAAATTTGTCGCCTGACTGTTTAAAGCTCGTTAGACTACGAT<br>ATTTTCCGCTTGTCGTAAGTTGTACAAGTAAATCAAGAATGATTTT<br>GTGATAGTACGGTTTAGACTGCCTGCTTTGCATGATTGCGGTGTCT<br>AGTTTGTTCATGGTTAGTTATCCTTAACTTGCAAAAAAATCAAGTT<br>AATAGTTAAAATTTTTCATCAAGTCATAAATAGAATTTTCTTCTAA<br>ATTTGCTGCTCTTTCTAATTCTTTAACCTTATCAAGTGTTAATTTAT<br>TCGGAGCTAATCTAATGCGATATAGAGCATTATATGTGATTCCCA<br>TATTCTTCGCTATCGCCTCATATCTTACCCCTGATTGTTTTAAAAT<br>CTCATCAAGTGGTTTATAAGTTTTACTCATTTTATCTCCTTTCTGAT<br>TTTTATGTTTTTCATTCTAACATTAACTTGATTTTTATGCAAGTAAT<br>AACTTTACTTTTTTGCAAGTTTTCTCTTGAAAGTAGTT |

**Supplementary Text**

Supplementary Note 1:

Cognate SSBs corresponding to each of the phage RecTs as follows:

λβ is found in an *Escherichia coli* phage. The selected strain *E. coli* K12 only has a single SSB protein annotated.

PapRecT is found in a *Pseudomonas aeruginosa* phage. The selected strain *P. aeruginosa* PAO1 only has a single SSB protein annotated.

MspRecT is found in a *Mycobacterium smegmatis* phage. The selected strain *M. smegmatis* mc(2)-155 has 3 annotated SSB proteins, however only one has the conserved tail motif (DDEPPF) similar to the SSBs selected for PaSSB and EcSSB so we selected that variant.

LrpRecT is found in a *Lactococcus reuteri* prophage. The selected strain *L. reuteri* MM4-1A has two annotated SSB proteins, however only one has the conserved tail motif (DDELPF) similar to the SSBs selected for PaSSB and EcSSB so we selected that variant.

Supplementary Note 2:
Metrics for calculating "SSB C7 compatibility", "Lower SSB C7 compatibility", and "Inhibitory SSB".

- An SSB was marked as "compatible" if it caused at least a 5x increase in recombination efficiency above the RecT alone, and was within 0.5x of the best performing SSB.
- An SSB was marked as having "lower SSB compatibility" if it caused between a 5x increase or 5x decrease in recombination efficiency, or contributed to an efficiency less then 0.5-fold of the best performing SSB.
- An SSB was marked as "inhibitory" if it caused a more than a 5x decrease in recombination efficiency.
- SSBs with the same C7 sequence were categorized together

Supplementary Note 3:
Calculation of estimated number of edited cells and library coverage for 1NNK and 5NNK libraries:

5x1NNK:
We performed a single transformation and plated half of the recovered cells (235 million) on 100µg/mL spectinomycin plates (117.5 million). We estimate that a single doubling occurred during the hour recovery (~58.75 million unique cells electroporated). The average efficiency of known single-amino acid changes at that position is 3.2% (V28P, K29I), giving an approximate number of 1.88 million edited cells. We assume complete coverage of the 105 amino acid variants (including stop codons).

5NNK:
We performed 30 parallel transformations and plated all of the recovered cells (6.75 billion) on 100ug/ml spectinomycin plates. We estimate that a single doubling occurred during the 60min recovery (3.375 billion unique cells electroporated). The average efficiency of 5AA mutations is 1.54% (RTNAR, NGTRF, Fig S12), giving an approximate number of 52.1 million edited cells.

The likelihood of sampling of a particular variant is given by: $1-(1-p)^{L}$ where $p_i$ represents the probability of each variant (i) and, L is the total number of edited cells. We calculate $p_i$ for each possible amino acid combination by assuming independence at each position, use the frequency of sampling each amino acid in the initial library (Fig S13), and take the product of the probabilities at each position.

For example: $p_{AGAAA} = p_{1A} \times p_{2G} \times p_{3A} \times p_{4A} \times p_{5A}$

To estimate total library coverage we then sum over all possible variants

n: $\sum_i^n 1-(1-p_i)^{L}$

Given L=52.1 million edited cells, our expected coverage is 3.95 million amino acid variants (including stop codons) or 96.7% of the total 4.08 million.

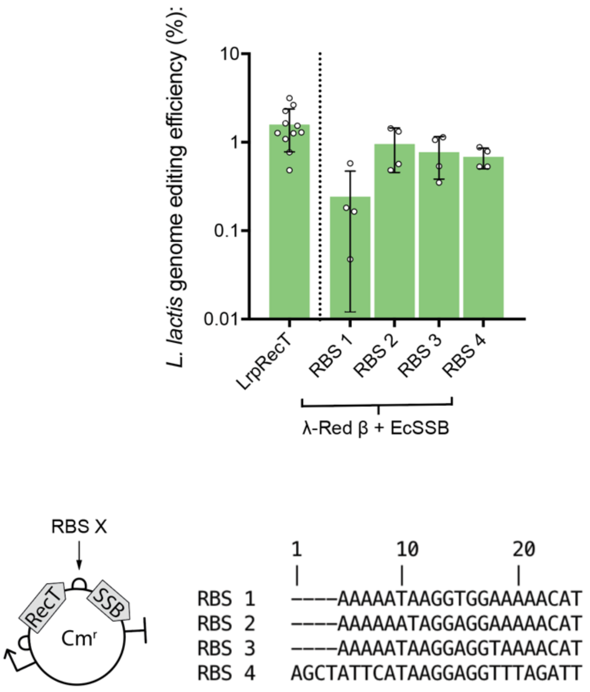


**Fig. S1.**
In *L. lactis*, the internal RBS sequence affected recombination efficiency using the bicistronic Redβ and EcSSB construct. RBS 2, which enabled the highest efficiency genome editing in this experiment was selected used in all other bicistronic constructs unless otherwise indicated.

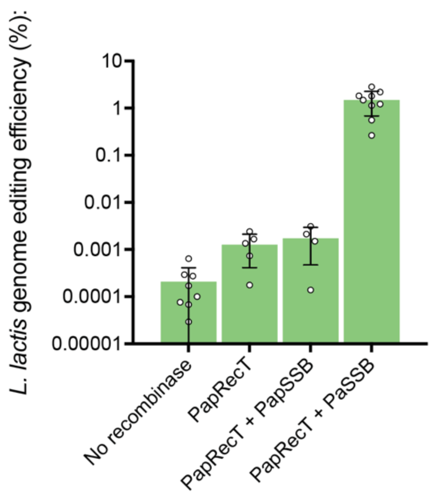


**Fig. S2.**

A number of phage operons containing a RecT protein also include a phage SSB protein, as is the case for the *P. aeruginosa* phage. However, co-expression of this phage SSB protein (PapSSB) along with the phage RecT (PapRecT) did not allow recovery of editing activity in *L. lactis*, unlike co-expression of a paired bacterial SSB (PaSSB).

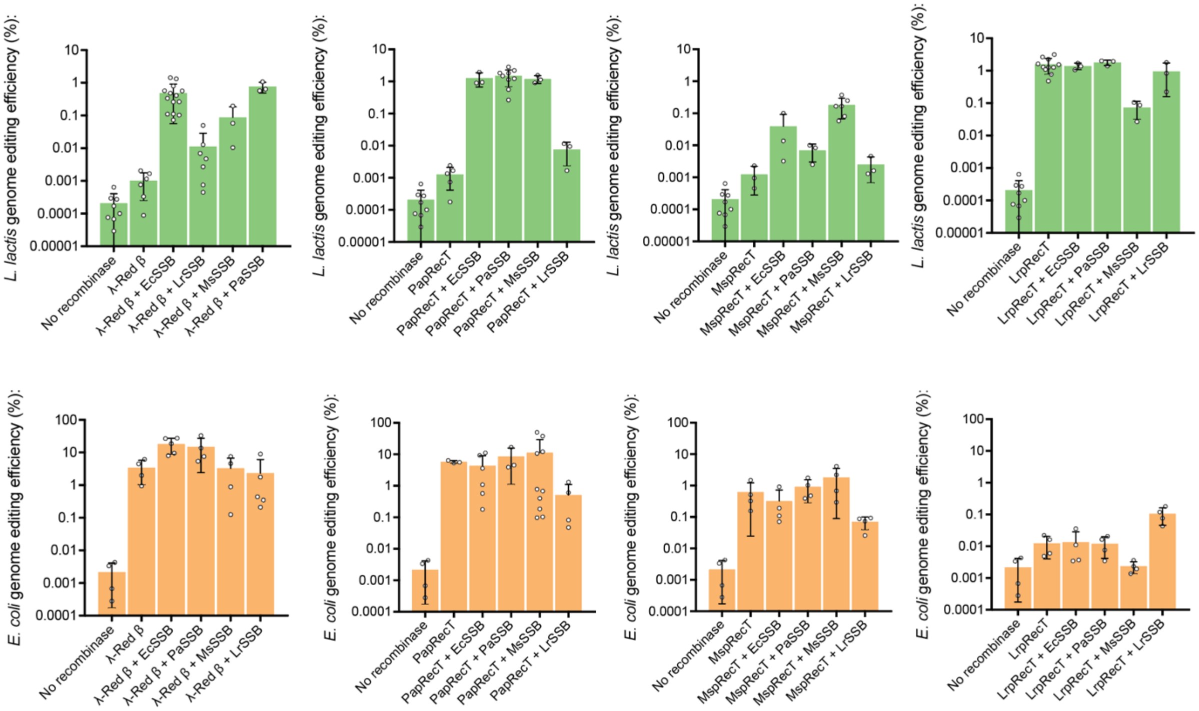


**Fig. S3.**

Bar graphs showing individual data points and SEM for mean values plotted in Fig 3a and Fig 3b.

We note there are four cases of SSB co-expression lowering RecT protein genome editing efficiencies. Specifically, LrSSB reduces the activity of MspRecT and PapRecT in *E. coli*, while MsSSB reduces the activity of LrpRecT in *E. coli* and *L. lactis*. We speculate that this may occur if the supplied SSB is close to compatible with the RecT protein but not fully compatible. In each of these cases, the 7 amino acid C-terminal tail of a compatible SSB is one amino acid different from that of the supplied SSB (Fig 3c).

We also observe two instances of SSB co-expression improving an otherwise active RecT protein (In *E. coli*: Redβ + EcSSB, and Redβ + PaSSB). This is unlikely to be caused simply by the reduction in toxicity (Fig S4), as some constructs have significantly reduced toxicities but similar genome editing activities (PapRecT with and without PaSSB).

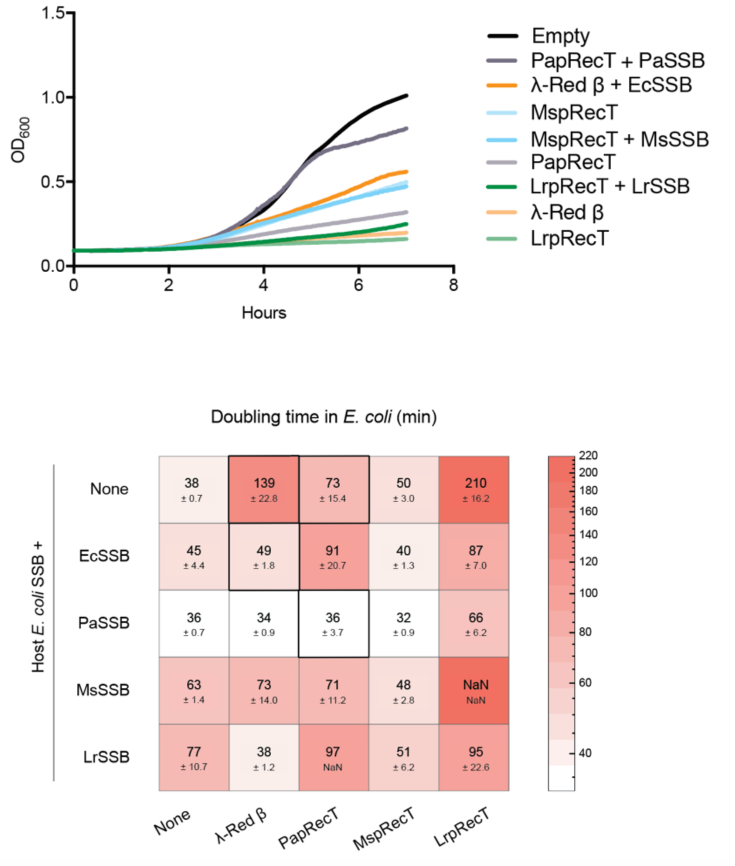


**Fig. S4.**

Doubling times in *E. coli* of constructs expressing RecT and SSB proteins reveal that co-expression of SSB can dramatically influence construct toxicity. Each of the constructs was inoculated into LB+Carb media containing the inducer arabinose (except for “Empty” which was inoculated into LB). The RecTs vary in toxicity, with Redβ showing considerable toxicity. The co-expression of SSBs reduces RecT toxicity in a number of cases, especially for PaSSB.

There are a number of constructs with both low toxicity and high activity (λβ + EcSSB, λβ + PaSSB, PapRecT + PaSSB) showing that there is no direct correlation between toxicity and activity.

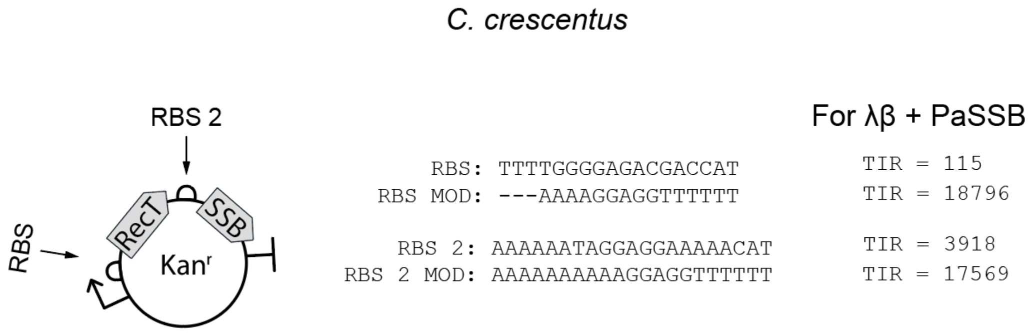


**Fig. S5.**
Using the Salis RBS calculator we designed RBSs conferring a greater translation rate in order to increase RecT and SSB expression for the Caulobacter constructs.

A. Espah Borujeni, A. S. Channarasappa, H. M. Salis, Translation rate is controlled by coupled trade-offs between site accessibility, selective RNA unfolding and sliding at upstream standby sites. Nucleic Acids Res. 42, 2646–2659 (2014).

H. M. Salis, E. A. Mirsky, C. A. Voigt, Automated design of synthetic ribosome binding sites to control protein expression. Nat. Biotechnol. 27, 946–50 (2009).

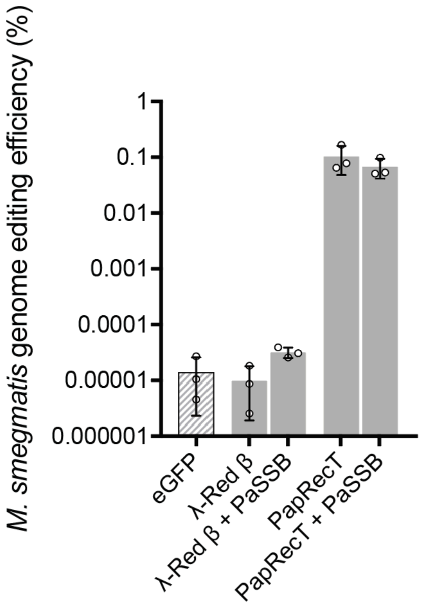


**Fig. S6.**

Recombineering using select RecTs and RecT-SSB pairs in *M. smegmatis*. Recombineering using Redβ + PaSSB is not substantially better than Redβ in *M. smegmatis*. PapRecT has basal activity in *M. smegmatis*, and PapRecT + PaSSB performs at approximately the same level.

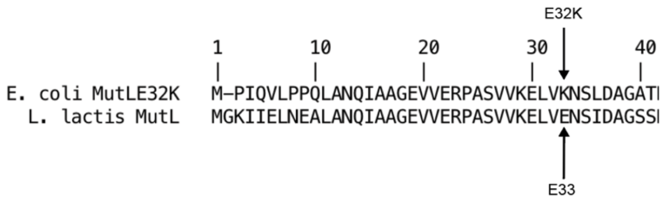


**Fig. S7.**

To generate a dominant negative *L. lactis* MutL, we aligned the host MutL protein with the dominant negative E. coli gene (MutL.E32K) and made the corresponding change in *L. lactis* (E33K).

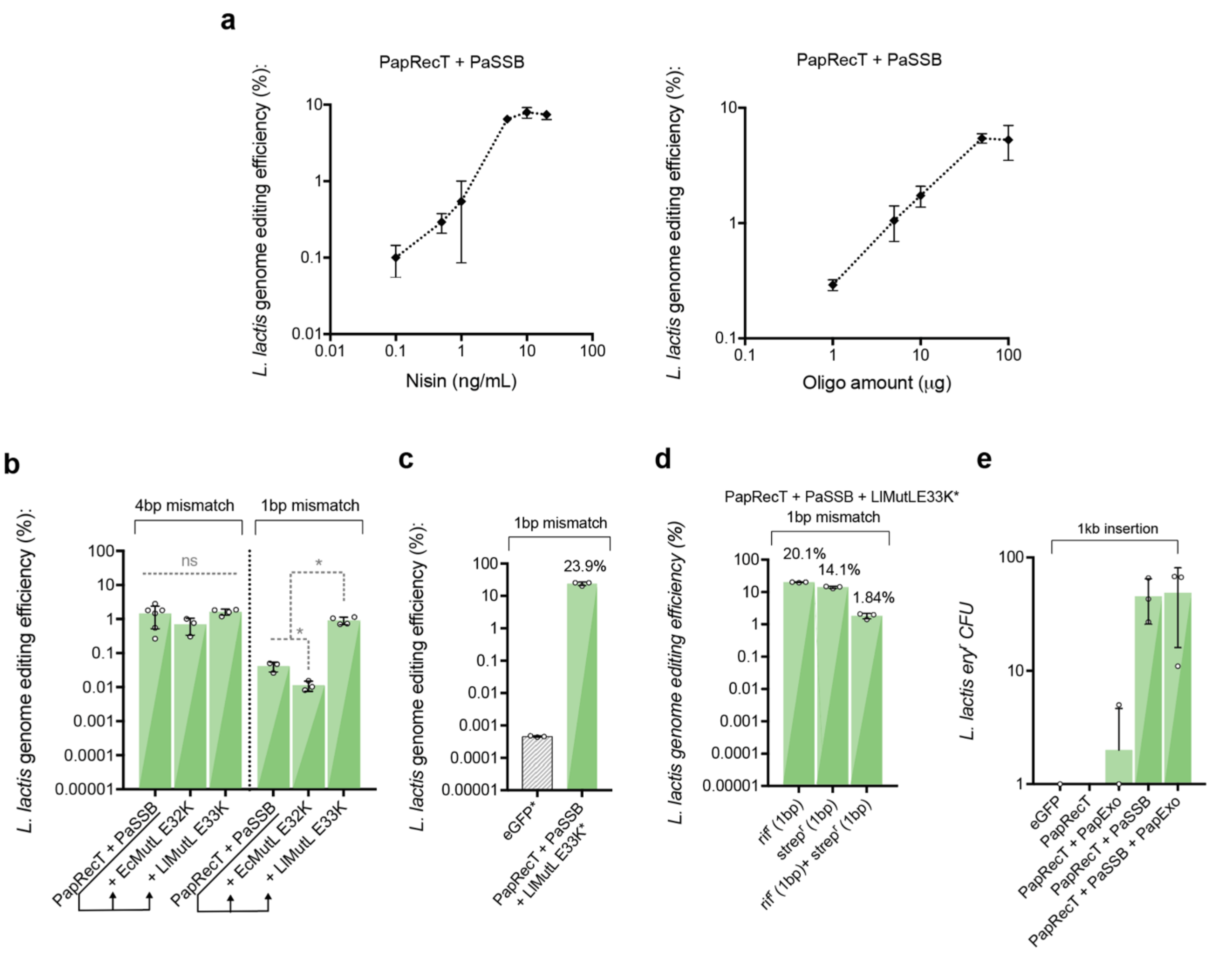


**Fig. S8.**

(a) In *L. lactis*, optimization of nisin concentration contributed to a significant improvement in editing efficiency for the PapRecT protein and the PaSSB protein construct. 10 ng/mL nisin was much more effective than 1 ng/mL nisin and resulted in an increase in editing efficiency improvement from 0.5% to 8%. The optimal oligo amount plateaued at 50 µg of DNA, which corresponds 21.4 µM in 80 µL. (b) Expression of the L. lactis MutL variant E33K allowed the efficient introduction of 1bp mismatches at similar efficiency to 4bp mismatches which evade MMR. (c) After optimization from (a,b), PapRecT + PaSSB + LlMutLE33K enabled ~20% editing efficiency at the Rif locus, and multiplexed editing (d). Co-expression of PapRecT + PaSSB enabled the efficient introduction of a 1kb selectable marker as dsDNA even without the addition of the cognate phage exonuclease. This also was observed for Redβ with EcSSB in *L. lactis* (Data not shown).

a

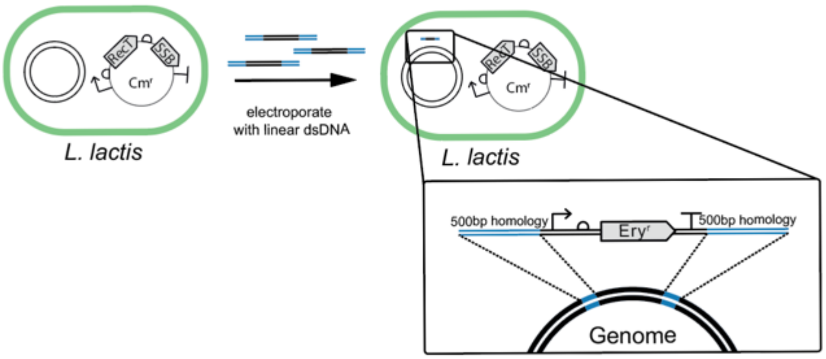


b

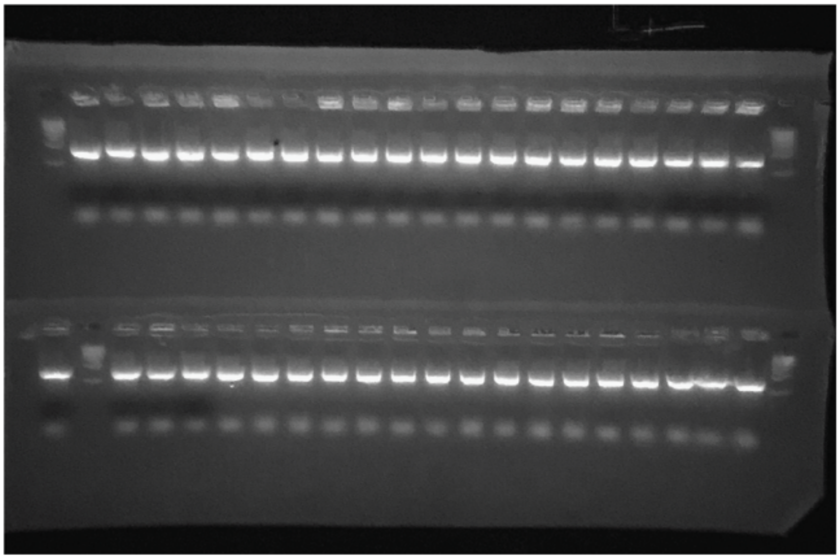

**Fig. S9.**
(A) Using linear DNA with 500bp homology arms we were able to generate gene knockins using PapRecT and PaSSB, which does not include an exonuclease (Fig S7).
(C), PCR amplification using primers within the genome outside of the homology arm and on the template verify insertion at the correct locus for all tested colonies. Correct band is 856bp long, ladder is NEB 1kb ladder.

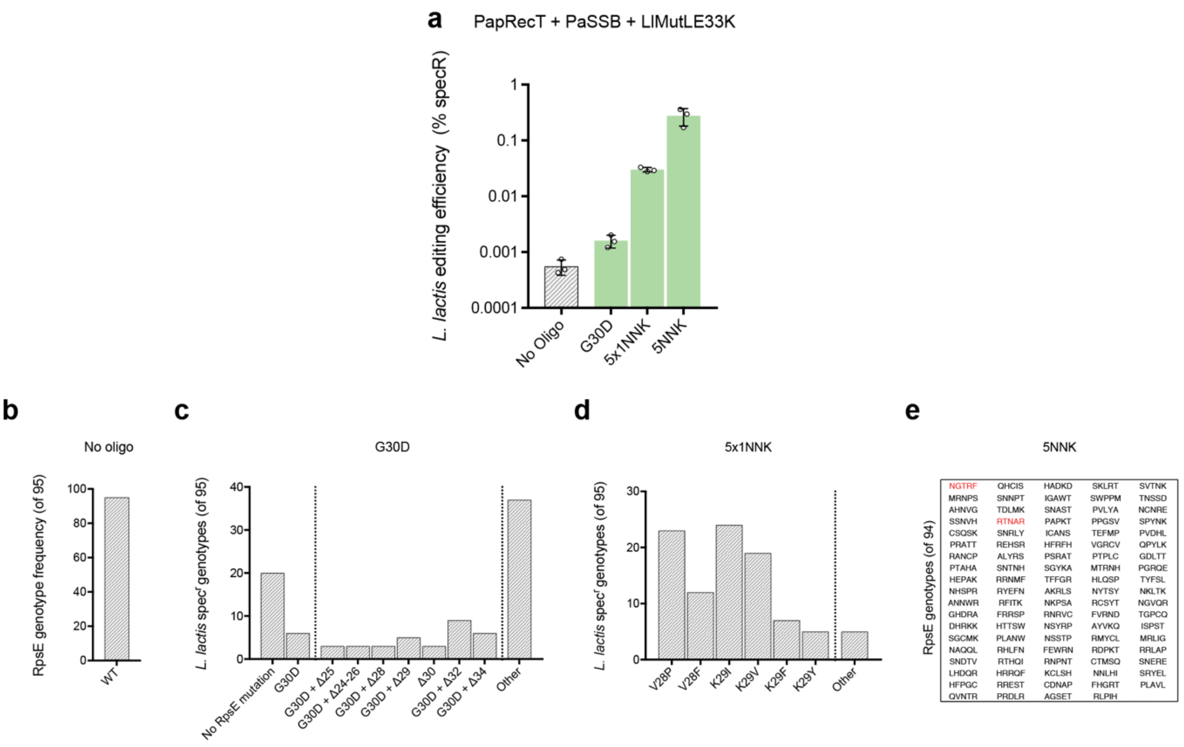


**Fig. S10.**
Preliminary experiment for spectinomycin resistance at *L. lactis* RpsE

(A), Efficiencies of spectinomycin resistance for different oligo designs measured in triplicate.

(B-E), Sanger sequencing of spectinomycin resistance mutants.

The amino acid conversion corresponding to a *E. coli* spectinomycin resistant mutant G27D (G30D) did not generate a resistant phenotype in *L. lactis*. The higher rate of resistant alleles using a G30D targeting oligo compared to background levels may be attributed to a rare occurrence of small deletions that occur during mutagenesis (b) which result in a resistant phenotype. To our knowledge this production of low levels of unintended deletions has not previously been described for recombineering, and may be due to oligonucleotide synthesis errors, or an aberrant effect of remaining mismatch repair proteins.

Sanger sequencing results from the 5x1NNK and 5NNK plates reflect allele mutations seen in the next-generation sequencing results

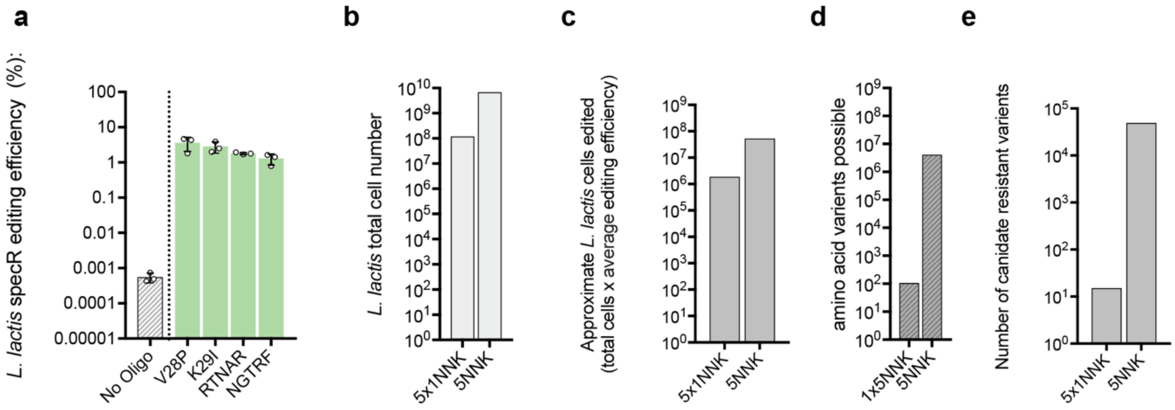


**Fig. S11.**

(A), Editing efficiency of select 1 amino acid and 5 amino acid variants.
(B), Total cell number after editing with the 5x1NNK oligo library (a single transformation), and the 5NNK oligo library (30 pooled transformations) targeting L. lactis RpsE.
(C). Calculation of expected number of edited cells using the average editing efficiency for 1AA changes and 5AA changes given in (A)
(D), Number of possible amino acid variants and stop codons in the two libraries (5 x 21 for 1x5NNK), (21^5 for 5NNK)

**Pre-selection 1NNK (WT reads removed)**

| | NNK1 | NNK2 | NNK3 | NNK4 | NNK5 |
|---|---|---|---|---|---|
| D | 0.007 | 0.007 | 0.009 | 0.008 | 0.007 |
| E | 0.006 | 0.005 | 0.004 | 0.005 | 0.005 |
| K | 0.009 | 0.786 | 0.008 | 0.009 | 0.009 |
| R | 0.013 | 0.015 | 0.015 | 0.013 | 0.802 |
| H | 0.009 | 0.011 | 0.011 | 0.009 | 0.011 |
| N | 0.013 | 0.017 | 0.014 | 0.015 | 0.014 |
| Q | 0.005 | 0.008 | 0.006 | 0.006 | 0.006 |
| Y | 0.009 | 0.010 | 0.011 | 0.010 | 0.010 |
| C | 0.006 | 0.006 | 0.007 | 0.006 | 0.006 |
| G | 0.007 | 0.006 | 0.797 | 0.798 | 0.006 |
| S | 0.019 | 0.020 | 0.019 | 0.018 | 0.020 |
| T | 0.016 | 0.020 | 0.017 | 0.016 | 0.017 |
| A | 0.012 | 0.010 | 0.009 | 0.011 | 0.009 |
| M | 0.007 | 0.008 | 0.007 | 0.007 | 0.006 |
| I | 0.011 | 0.011 | 0.010 | 0.010 | 0.010 |
| L | 0.018 | 0.019 | 0.017 | 0.017 | 0.019 |
| V | 0.803 | 0.008 | 0.009 | 0.010 | 0.010 |
| F | 0.007 | 0.007 | 0.008 | 0.007 | 0.008 |
| W | 0.004 | 0.004 | 0.004 | 0.004 | 0.003 |
| P | 0.013 | 0.016 | 0.014 | 0.013 | 0.014 |
| * | 0.007 | 0.007 | 0.006 | 0.008 | 0.006 |

**Pre-selection 5NNK (WT removed)**

| | NNK1 | NNK2 | NNK3 | NNK4 | NNK5 |
|---|---|---|---|---|---|
| D | 0.035 | 0.034 | 0.037 | 0.036 | 0.036 |
| E | 0.024 | 0.023 | 0.024 | 0.023 | 0.023 |
| K | 0.042 | 0.044 | 0.041 | 0.041 | 0.040 |
| R | 0.064 | 0.073 | 0.064 | 0.067 | 0.074 |
| H | 0.045 | 0.045 | 0.048 | 0.048 | 0.047 |
| N | 0.064 | 0.060 | 0.065 | 0.068 | 0.066 |
| Q | 0.030 | 0.032 | 0.032 | 0.030 | 0.030 |
| Y | 0.046 | 0.044 | 0.047 | 0.049 | 0.050 |
| C | 0.025 | 0.028 | 0.025 | 0.027 | 0.029 |
| G | 0.030 | 0.033 | 0.033 | 0.034 | 0.032 |
| S | 0.090 | 0.090 | 0.087 | 0.089 | 0.090 |
| T | 0.088 | 0.083 | 0.083 | 0.082 | 0.079 |
| A | 0.053 | 0.051 | 0.051 | 0.047 | 0.046 |
| M | 0.033 | 0.033 | 0.032 | 0.031 | 0.032 |
| I | 0.043 | 0.043 | 0.045 | 0.048 | 0.047 |
| L | 0.083 | 0.086 | 0.087 | 0.084 | 0.085 |
| V | 0.048 | 0.043 | 0.045 | 0.045 | 0.043 |
| F | 0.033 | 0.033 | 0.034 | 0.035 | 0.037 |
| W | 0.017 | 0.019 | 0.016 | 0.017 | 0.018 |
| P | 0.074 | 0.074 | 0.073 | 0.067 | 0.065 |
| * | 0.030 | 0.030 | 0.031 | 0.030 | 0.032 |

**Fig. S12.**

Normalized frequencies of next-generation sequencing reads in the 1x5NNK library, as well as the 5NNK library after editing and 1hr of recovery, before selection with spectinomycin.

**a**

| 5NNK | Genotype | Resistant 100ug/ml |
|---|---|---|
| 3 AA mutant | VMNRR | Yes |
| 2 of 3 | VMNGR | Yes |
| 2 of 3 | VMGRR | No |
| 2 of 3 | VKNRR | Yes |
| 1 of 3 | VMGGR | No |
| 1 of 3 | VKNGR | No |
| 1 of 3 | VKGRR | No |

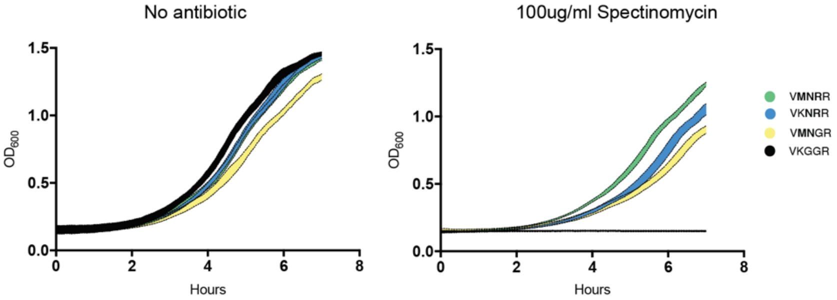


**b**

| 5NNK | Genotype | Resistant 100ug/ml |
|---|---|---|
| Most enriched | SNNRK | Yes |
| 3 AA mutant | RKNNR | Yes |
| 2 of 3 | RKNGR | Yes |
| 2 of 3 | RKGNR | No |
| 2 of 3 | VKNNR | Yes |
| 1 of 3 | RKGGR | No |
| 1 of 3 | VKNGR | No |
| 1 of 3 | VKGNR | No |

| 1NNK | Genotype | Resistant 100ug/ml |
|---|---|---|
| Most enriched | PKGGR | Yes |
| 2nd most enriched | VIGGR | Yes |

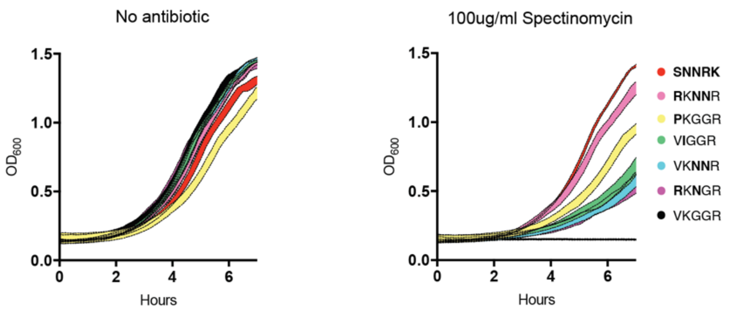


**Fig. S13.**

Doubling times of additional combination mutants and their resistant constituent variants (a, VMNRR) (b, RKNNR, as well at the most enriched variant in the entire pool SNNRK).

The triple mutant VMNRR is more resistant than its two constituent two double mutants (VKNRR, and VMNGR). Neither the remaining double mutant VMGRR nor any of the single mutants grew on 100 µg/mL Spec plates.

SNNRK grew faster in the presence of antibiotic than any of the other variants, and had less of a fitness defect than PKGGR even though all 5 positions are mutated from wildtype. Again, the triple mutant RKNNR was more resistant than its constituent double mutants and none of the signal mutants grew on 100 µg/mL Spec plates.

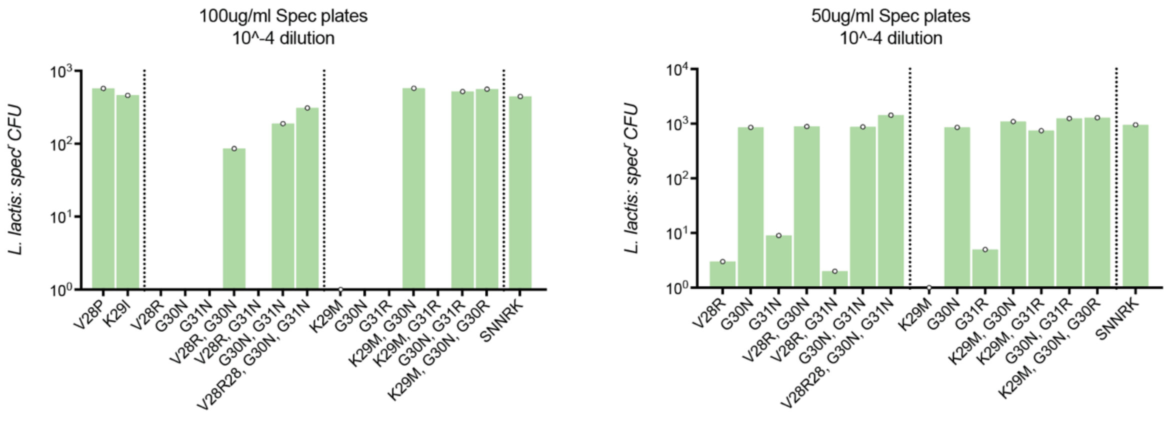


**Fig. S14.**

Validation of resistance for two genotypes (VMNRR, RKNNR) after two days of growth on 100 µg/mL spectinomycin plates or 50 µg/mL spectinomycin plates.

One of most enriched alleles that appeared during combinatorial mutagenesis is G30N. This mutation alone did not allow colony formation after 2 days on 100 µg/mL Spec plates, but did on 50 µg/mL Spec plates, indicating that weaker selection pressure may provide additional paths to highly resistant phenotypes.

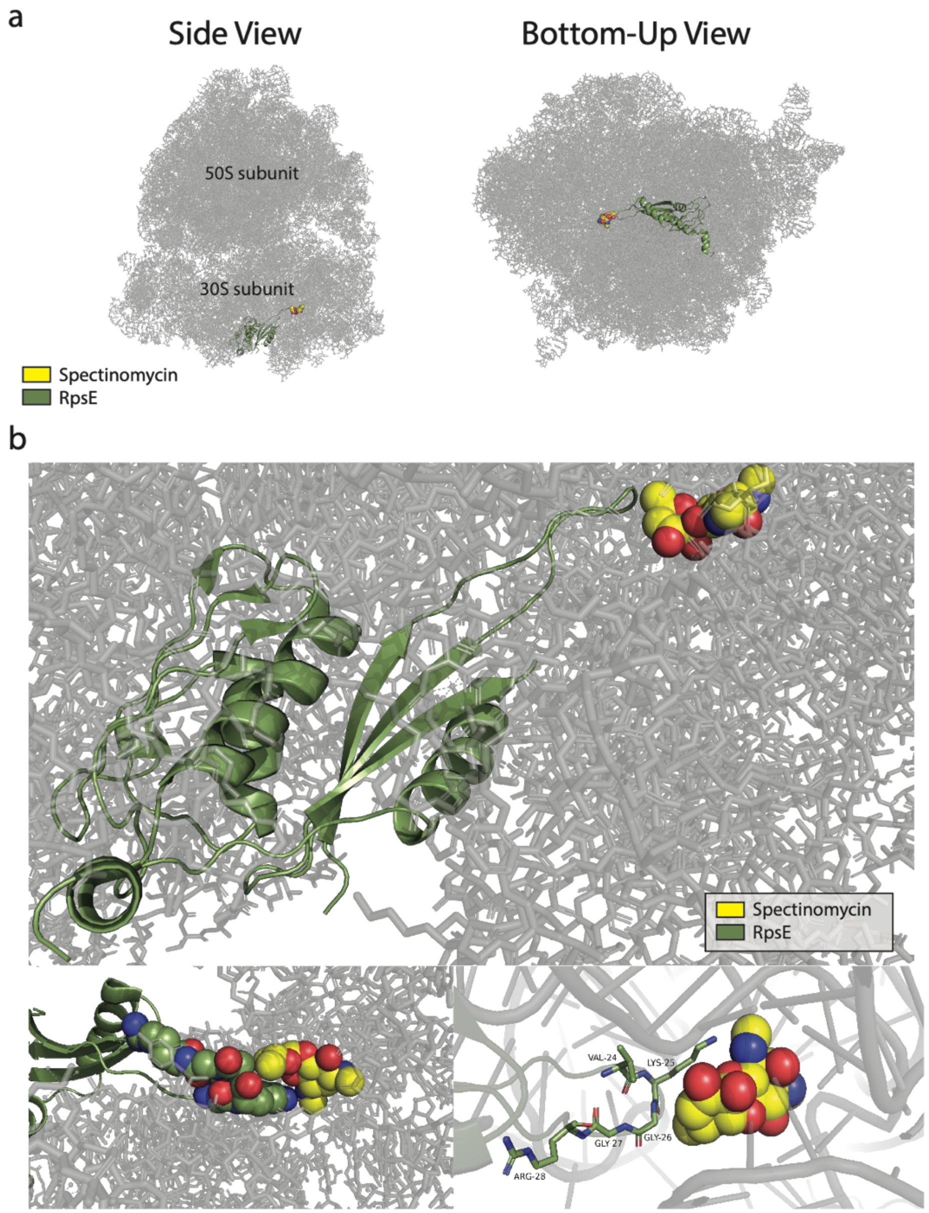


**Fig S15.** Structure Supplement. (a) Side (left) and bottom-up (right) views of the *E. coli* ribosome bound to spectinomycin (PDB ID: 4V56). Spectinomycin is shown in yellow spheres, RpsE (the protein product of ribosomal protein S5) in green ribbon, and the rest of the RNA and protein components of the ribosome in gray, partially transparent sticks. (b) The bottom-up view is magnified to show the interaction between RpsE and Spectinomycin. Three different views are shown: (top) the full RpsE protein is visible in green ribbon, (bottom left) the loop that contacts spectinomycin is focused on with the five varied residues shown in green spheres, and (bottom right) this same loop is focused on with residues labeled and shown in green sticks.

| Figure | Statistical Test | Multiple comparisons |
|---|---|---|
| Fig 1g. | Welch's two tailed t-test of Log-transformed data; P value < .05 | |
| Fig 1h. | Welch's two tailed t-test of Log-transformed data; P value < .05 | |
| Fig 2c. | Ordinary one-way ANOVA of Log-transformed data; P value < .05 | Holm-Sidak |
| Fig 2d. | Ordinary one-way ANOVA of Log-transformed data; P value < .05 | Holm-Sidak |
| Fig 2e. | Ordinary one-way ANOVA of Log-transformed data; P value < .05 | Holm-Sidak |
| Fig 2f. | Ordinary one-way ANOVA of Log-transformed data; P value < .05 | Holm-Sidak |
| Fig 3d. | Ordinary one-way ANOVA of Log-transformed data; P value < .05 | Holm-Sidak |
| Fig 3e. | Ordinary one-way ANOVA of Log-transformed data; P value < .05 | Holm-Sidak |
| Fig 4a. | Welch's two tailed t-test of Log-transformed data; P value < .05 | |
| Fig 4b. | Welch's two tailed t-test of Log-transformed data; P value < .05 | |

**Table S1. Statistical tests**
